# A phylogenetic estimate of canine retrotransposition rates based on genome assembly comparisons

**DOI:** 10.1101/2025.09.10.675418

**Authors:** Matthew S. Blacksmith, Anthony K. Nguyen, John V. Moran, Jeffrey M. Kidd

## Abstract

**Background:** Due to their history of domestication and breed formation, dogs are a powerful system for studying the phenotypic impact of genetic variation. Comparison of canine genome assemblies show that retrotransposons, mobile elements that mobilize via an RNA intermediate, are a major contributor to canine genetic diversity with an eightfold and 17-fold increase of LINE-1 and SINE differences, respectively, in dogs relative to that found among humans. The frequency of dimorphic retrotransposon insertions among dogs suggests these elements have mobilized at a high rate over recent canine evolution. However, the rate at which new insertions arise has yet to be determined.

**Results:** We aligned a collection of genome assemblies derived from four breed dogs, a Dingo, and an American grey wolf to a Greenland grey wolf to identify dimorphic LINE-1 and SINEC insertions. Across our panel of seven canine assemblies, we identified and characterized 7,428 dimorphic LINE-1s and 51,572 dimorphic SINECs. Each assembly differs from the Greenland wolf genome by an average of 3,497 LINE-1s and 25,558 SINECs. Analysis of allele sharing among samples recapitulates known relationships and reveals substantial within-breed variation. Calibrating estimates using a previously estimated single nucleotide mutation rate of 4.5x10^-9^ per base pair per generation, we estimate that new LINE-1 and SINEC and insertions have occurred at a rate of 1/184 and 1/22 births over recent canine evolution. These estimates are largely consistent across assemblies and breeds.

**Conclusions:** Our phylogenetic estimate of SINEC retrotransposition in canines is approximately twice as large as that estimated for *Alu* elements in humans, while the canine LINE-1 insertion rate is within the range of human estimates. These data suggest that although SINEC has been an outsized driver of canine genome evolution, the striking levels of canine LINE-1 and SINEC dimorphism mainly reflect high levels of long-standing genetic variation.

## Background

Dogs, *Canis lupus familiaris*, have been a human companion for over ten thousand years. Recent findings using ancient DNA show that dogs were domesticated from a now-extinct Eurasian gray wolf lineage approximately 10-40 thousand years ago (1–4). Since domestication, dogs have been subject to extensive selection by humans based on their behavior, size, and morphology (5–9). Selection became more extreme with the formation of modern breeds in the late 19^th^ century (10). This active process has seen the creation of over 300 recognized breeds worldwide that show over an order of magnitude of size difference (e.g., Great Dane: 80kg, Chihuahua: 0.5kg) (11).

The natural genetic variation found in dogs, along with the unique structure of breeds, has made dogs a powerful system for mapping the molecular basis of traits and studying the impact of selection and population bottlenecks (9, 12–14). The potential of dogs as a model system has led to the creation of a robust canine genomics research community and catalyzed the development of multiple genomic resources including numerous genome assemblies and myriad short read datasets (15, 16). One striking finding from the analysis of these resources is the high level of retrotransposon differences found among dogs. Retrotransposons are a type of mobile element that propagates throughout the genome using a “copy and paste” mechanism via an RNA intermediate (reviewed in (17)). Retrotransposons are further subdivided into two classes, those that contain long terminal repeats (LTRs) and those that do not. Non-LTR retrotransposons, which include Long INterspersed Elements (LINEs) and Short INterspersed Elements (SINEs), are highly prevalent in the canine genome (18).

Comparisons between the initial dog reference genome, derived from a Boxer breed dog named Tasha, and low-coverage survey sequencing from a Poodle breed dog, named Shadow, revealed over 10,000 SINEs with presence-absence differences (known as dimorphism) between the two assemblies (18, 19). Comparison of genome assemblies from a Boxer and a Great Dane identified over 16,000 SINECs and 1,100 LINE-1s that differed between the assemblies (20). Subsequent analyses demonstrated that a full-length LINE-1 cloned from the Great Dane genome was active in a cell culture based retrotransposition assay (20, 21). The cloned canine LINE-1 sequence was also capable of driving SINEC retrotransposition in cell culture (20, 22). Recent years have also seen an explosion in canine genome comparisons, further confirming the outsized contribution of retrotransposon sequences to canine genetic variation (16, 20).

LINE-1, the family of LINEs currently active in mammals, including canines, is approximately 6 kb in length and encodes two proteins (23, 24). ORF1p contains RNA binding and nucleic acid chaperone activities, while ORF2p contains domains responsible for nicking genomic DNA (endonuclease) and copying the element-encoded RNA into DNA (reverse transcriptase) (25–30). LINE-1 retrotransposition begins when LINE-1 is transcribed by RNA polymerase II (Pol II), using the internal promotor present in the LINE-1 5’ UTR (31, 32). LINE-1s also possess a polyadenylation signal and are dependent upon a 3’ poly(A) tail to fully proceed through the retrotransposition cycle (33–35). Following transcription, LINE-1 RNA is exported to the cytoplasm where the bicistronic RNA is translated by an unconventional mechanism (36). In humans, ORF1p is translated to a higher degree and is reliably present in cytosolic fractions (28, 37). By contrast, ORF2p is produced in lower abundance and co-translationally associates with the 3’ poly(A) tail of the LINE-1 RNA that encodes it (35, 38, 39). Once bound by both ORF1p and ORF2p, the ribonucleoprotein particle is imported back into the nucleus to undergo a process termed target primed reverse transcription (TPRT) (21, 29, 40–42). During TPRT, the ORF2p endonuclease activity creates a nick in genomic DNA at a new locus, with a preference for the ORF2p canonical cleavage site of 5’-TTTTT/AA (29, 43, 44). Subsequently, the ORF2p reverse transcriptase activity is used to generate a DNA copy of the LINE-1 RNA which, following second strand synthesis, results in the insertion of a copy of the LINE-1 RNA into a new genomic locus (21, 30, 45). LINE-1 insertions are often severely 5’ truncated or contain other mutations that render them inactive (46–48).

LINE-1 mediated insertions possess several hallmarks which can be used to characterize the insertion (reviewed in (17) and (49)). The first, mentioned above, is the presence of the canonical endonuclease cleavage site, 5’TTTTT/AA, at the site of insertion (21, 29, 43, 44) . Second, is the presence of a 3’ poly(A) sequence which is placed by a poly(A) polymerase post-transcriptionally and is required for the binding of ORF2p (33, 35). Third, is the presence of target site duplications (TSDs), which arise during TPRT and flank the insertion (43, 49, 50). Finally, while not present in all LINE-1 insertions, 3’ transductions are frequently identified in LINE-1 datasets (21, 33, 51). LINE-1 has a weak termination signal and consequently, RNA polymerase II may read through the LINE-1 termination signal into downstream sequence (21, 33, 51, 52). This process creates a transcript containing LINE-1 and downstream genomic sequences; the genomic sequence can then be used to infer the identify of a parent element associated with an insertion (20, 51, 53–56).

In addition to mobilizing the transcripts that encoded them (known as *cis*-preference (39, 41)), the LINE-1 encoded proteins are able to mobilize other RNAs (57), the primary example of which are SINEs. SINEs do not encode proteins, but instead rely upon LINE-1 ORF2p to mobilize (22). A family of SINEs, termed SINEC, is present throughout *Carnivora* and is abundant in the canine genome (18, 19, 58). SINEC sequences are transcribed by RNA polymerase III (Pol III) and contain a 5’ ‘head’ region derived from tRNA-lysine, a segment of (CT)_N_ repeats of variable lengths, and an A-rich tail (58–60). It is hypothesized that SINEC may retrotranspose via two similar mechanisms: one that utilizes a cellular poly(A) polymerase to generate a poly(A) tail, and one that uses the existing poly(A) sequence as a poly(A) tail (61). LINE-1 encoded proteins can also mobilize other cellular RNAs, leading to the production of retrocopies, also known as processed pseudogenes or retrogenes (41, 57).

LINE-1 mediated insertions contribute to, or are associated with, numerous canine phenotypes and diseases (7). Perhaps the most well-known are the multiple functional retrocopies of the *FGF4* gene that cause a chondrodysplasia-like short-legged phenotype in breeds such as the Pembroke Welsh Corgi, Basset Hound, Dachshund, and Nova Scotia Duck Tolling Retrievers (62, 63). LINE-1 insertions are also implicated in progressive retinal atrophy in Swedish Vallhunds, Duchene-like muscular dystrophy in Pembroke Welsh Corgis, and brachycephaly in several breeds (64–66). Conditions such as polyneuropathy are caused by a SINEC insertion into *RAB3GAP1* in Alaskan Huskies (67), and variability in a SINEC insertion causes the merle coat color pattern found in some breeds (68, 69). Importantly, a recent *de novo* LINE-1 insertion into the *DMD* gene has been identified in a Border Collie, demonstrating that LINE-1 continues to actively mobilize in the canine genome (70).

The frequency of dimorphic LINE-1 and SINEC insertions among dogs may reflect both a high mobilization rate over recent canine evolution and the assortment of standing genetic variation that arose in ancestral wolf populations (6, 7). Using LINE-1 and SINEC dimorphic elements identified by comparison of canine genome assemblies, we produce a phylogenetic estimate of the rate of LINE-1 and SINEC retrotransposition over recent canine evolution that is calibrated using a previously estimated single nucleotide variant (SNV) mutation rate. Our analysis confirms the high degree of LINE-1 and SINEC dimorphism in canines, including between individuals of the same breed. We estimate that new LINE-1 and SINEC insertions arise at a rate of 1/184 and 1/22 births, respectively, in canines.

## Methods

### Identification of dimorphic loci

Structural variants that correspond to dimorphic LINE-1 or SINEC elements were identified from seven published canine genome assemblies, including two German Shepherd Dogs (GSD1, GSD2), a Boxer (BOX), a Great Dane (GDN), a Dingo (DNG), and two grey wolves (A_WOLF, G_WOLF) (20, 71–76) (Table 1, Supplemental Data S1). For ease of description, we refer to each assembly based on the shortened name given in Table 1 and Supplemental Data S1. Variants were identified by aligning each genome to that of the Greenland wolf (G_WOLF), which was assembled using highly accurate PacBio HiFi reads and serves as an outgroup to domestic dogs.

**Table 1:** Sample demographics and accession information.

| **Sample Key\*** | **Breed/Species** | **Sample Name** | **Segment Count** | **Assembly Name** | **Accession** |
| --- | --- | --- | --- | --- | --- |
| **G_WOLF** | Greenland wolf | mCanlor1.2 | 205 | mCanlor1.2 | GCA_9053 |
| **A_WOLF** | American wolf | Clu | 111 | GCA_034620435.1_Clu-1 | GCA_0346 |
| **GSD1** | German Shepherd Dog | Mischka | 402 | UU_Cfam_GSD_1.0/canFam4 | GCA_0111 |
| **GSD2** | German Shepherd Dog | Nala | 632 | CanFam_GSD | GCA_0086 |
| **BOX** | Boxer | Tasha | 1,067 | Dog10K_Boxer_Tasha_1.0/canFam6 | GCA_0000 |
| **GDN** | Great Dane | Zoey | 1,036 | UMICH_Zoey_3.1/canFam5 | GCA_0054 |
| **DNG** | Dingo | Sandy | 107 | CanFam_DDS | GCA_0032 |

\*Sample Key refers to the name of the sample used in figures and legends throughout the paper.
\*\*A segment is defined as any autosomal or chromosome X sequence that is uninterrupted by nucleotides of unresolved sequence (N). A complete telomere to telomere assembly would have 39 segments.

Each of the six assemblies were aligned as queries using the G_WOLF assembly as the target using minimap2 (version 2.26) (77) with options -c -x asm5 --cs. The resulting alignment files in paf format were then sorted (sort -k6,6 -k8,8n) and processed using the paftools.js call utility with default parameters to create a listing of variants identified from each pairwise comparison. SNVs and structural variants (SVs) at least 50 bp in size, were extracted from the variant list and filtered to include only variants on the autosomes (chr1-chr38) or X chromosome and where the query and target sequences occur on the same chromosome.

Next, a list of exclusion regions was aggregated, including segmental duplications identified by BISER (78) and/or fastCN (79), as well as genome assembly gaps. Duplication coordinates from G_WOLF, GSD1, GSD2, BOX, GDN, and DNG were obtained from Nguyen, Blacksmith, and Kidd, 2024 (80). Duplications were identified in A_WOLF following the same procedure (80). Because Illumina reads of A_WOLF are not directly available, we used Illumina reads from the wolf parent of A_WOLF. Exclusion regions were defined in both the query and target. Any SV within 100 bp of an excluded region was removed from the dataset using bedtools window with options -w 100 -v (version 2.26.0) (81). SVs were then split into “insertions” and “deletions” before being further divided into candidate LINE-1 and SINEC elements based on comparison with RepeatMasker annotations (version 4.0.7 Database: dc20170127-rb20170127) using bedtools intersect with options -wa -f 0.7. The -f option requires that 70% of each SV is represented in the corresponding RepeatMasker file (82). LINEs belonging to the HAL family were excluded from analyses (83). This criterion is designed to distinguish structural variants that correspond to dimorphic mobile element insertions from other structural variants that contain mobile element sequences. In our informatics pipeline, we defined insertions as variants present in an analyzed sample but absent from G_WOLF and deletions as variants present in G_WOLF but absent from the sample being compared. In reality, dimorphic mobile element insertions represent two states: (***1***) the ancestral “empty site” allele without the element, and (***2***) the derived or “filled site” allele that contains the mobile element. The mobile element content of the filled site allele identified for each variant was initially determined based on the annotation of the appropriate genome assembly. Additionally, sequence annotations were determined by running RepeatMasker directly on the extracted sequence corresponding to each filled site. Sequence coordinates and annotations are available in Additional_File1. Annotations were determined based on RepeatMasker (version 4.0.7 Database: dc20170127-rb20170127). The sample standard deviation for the number of SINECs present across the dataset was calculated using numpy.std(data,ddof=1) (v1.20.3).

### Identifying the hallmarks of retrotransposition in SINEC and LINE-1 loci

The hallmarks of LINE-1 retrotransposition (i.e., TSDs, 3’ poly(A) sequences, and endonuclease cleavage sites) in dimorphic mobile element containing loci were identified by comparing the filled (SINEC or LINE-1 present) and empty (SINEC or LINE-1 absent) genomic DNA sequences. For each locus, the filled and empty site sequence, including 1,000 bp of flanking sequence in each direction, were extracted with samtools faidx (v1.21) (84) and aligned with AGE (v0.4) (85). AGE is an alignment software designed for the analysis of structural variants and can identify the presence of matching sequences located at rearrangement breakpoints. Because AGE can only accurately resolve one structural variant per alignment, if the variant identified by AGE is more than 150 bp away from the coordinate originally identified from the minimap2 alignment, the AGE processing was repeated using a reduced flanking size of 500 bp. If necessary, this process was additionally repeated two times using flanking sizes of 250 bp and 125 bp, respectively. If consistent coordinates could not be identified, the locus was excluded from further processing. Several other filters were also introduced, including removal of loci which are: (***1***) less than 50 bp in length after AGE processing, or (***2***) where the number of inserted base pairs that are not called as TSD is below 30. The presence of deletions in existing LINE-1 sequences gave rise to systemic LINE-1 false positives in this dataset. Two additional filters were used to remove these loci from further analysis. First, for each inserted sequence identified by AGE, RepeatMasker was used to directly assess the LINE-1 content of the inserted sequence. If less than 70% of the sequence was identified as LINE-1 sequence using this method, the locus was discarded from further processing. Second, utilizing the original whole genome RepeatMasker annotation, loci which have a RepeatMasker identified segment extending beyond the inserted sequence as identified by AGE were checked for the presence of a TSD greater than or equal to 10 bp. If less than a 10 bp TSD was identified, these loci were discarded from further analyses. For the remaining SINEC and LINE-1 loci with concordant coordinates identified by AGE and minimap2, over 99.8% of the TSDs identified by AGE show perfect identity among the upstream TSD, downstream TSD, and the corresponding empty site. TSDs 10 bp or longer were considered “high confidence.”

To identify 3’ poly(A) tails, all sequence between TSDs, or the end of the insertion if no TSD is present, were searched for homopolymers of “A” or “T” depending on element orientation. Element orientation was identified by bedtools intersecting the query sequence coordinates mentioned above with SINECs or LINE-1s identified via RepeatMasker (81, 82). RepeatMasker segments smaller than 20 bp were not used for orientation detection. If multiple RepeatMasker segments are detected in a single locus and all segments are in the same orientation, the locus is fully processed; otherwise, the locus is retained but the poly(A) tails are not identified. If no RepeatMasker segments meet this requirement the locus is removed from processing. Homopolymers separated by 5 or fewer bases from the TSD (or insertion boundary if no TSD is present) were classified as poly(A) tails. If two or more homopolymers meet these criteria, the longer is chosen as the poly(A) tail. Only identified segments that are 10 bp or longer are retained as poly(A) tails.

Additionally, we implemented a relaxed hallmark identification pipeline to identify variants with shorter target site duplications and/or degraded poly(A) tails. We first reduced the minimum target site duplication length to seven bp instead of ten. Then, if the target site duplication was at least five bp in length, we extracted the 30 bp adjacent to the target site duplication present at the 3’ end of the insertion. Otherwise, the terminal 30 bases of the insertion were extracted. The extracted bases were scanned 15 bases at a time; if at least one window was found with 10bp of “A” content the locus was determined to possess a poly(A) tail.

Endonuclease cleavage sites were only identified in loci possessing a TSD of 10 bp or longer and a defined element orientation. Presuming that a locus is in the forward orientation relative to the reference, the endonuclease cleavage site consists of the two nucleotides upstream of the 5’TSD and the first 5 nucleotides of the TSD. All endonuclease cleavage sites are reported on the minus strand and thus should correlate with the known “5’-TTTTT/AA” cut site (43, 44). Sequence logoplots were created using logomaker (v0.8) (86). Other figures associated with this manuscript were generated using numpy (v1.20.3) (87), pandas (v1.3.4) (88), matplotlib (v3.4.3) (89), and scipy (v1.7.1) (90).

### Visualizing loci with miropeats

Filled and empty site sequences corresponding to dimorphic elements were extracted with samtools faidx (84). Each sequence is then investigated for repeats using RepeatMasker with the option --species dog (82). From there, a baseline miropeats image is generated using miropeats with options -onlyinter -s 200 (v2.02) (91). The miropeats image is then annotated to display RepeatMasker information.

### Identifying SINEC and LINE-1 subfamilies

To identify which subfamily each identified variant belongs to, coordinates corresponding to the sequence between target site duplications, or the full locus if no TSD is present, were intersected with all SINEC loci identified in genome-wide RepeatMasker annotation using bedtools intersect (81, 82). As in 3’ poly(A) identification, the intersected locus must be at least 20 bp in length. If multiple different subfamilies were identified, the locus is not uniquely assigned to a subfamily.

RepeatMasker reports 21 different SINEC subfamilies. This includes SINEC_Cf, a consensus sequence used by RepeatMasker that is not present in Repbase. Because the tRNA-derived head region of the SINEC_Cf consensus is identical to the SINEC2A1_Cf sequence, with the only differences being the length of the internal (CT)_N_ repeat, the length of the encoded poly(A) tail, and the presence of an extra T in the 3’ end of the element, we combined RepeatMasker annotations into a single subfamily we refer to as SINEC_Cf/2A1. Similarly, LINE-1 subfamilies were identified using the above methods.

### Identifying shared dimorphic SINEC and LINE-1 loci

To identify which SINEC and LINE-1 loci were shared across the dataset, we first identified autosomal regions of the G_WOLF genome that were covered by one query contig in each comparison, as reported by paftools.js call. Due to differences in X chromosome assembly quality, locus sharing analysis was limited to the autosomes.

The callable regions were further filtered to remove segments corresponding to duplications and gaps in the G_WOLF assembly as described above. We then created a bed file containing the intervals that were not callable in at least one sample and removed any dimorphic SINEC locus that intersected with the non-callable regions.

Remaining SINEC loci were then aggregated, sorted (with utilities sort -k 1,1 -k 2,2n -k 3,3n), and merged with bedtools merge -d 100 -c 7,4 -o collapse, collapse. These parameters merged any loci within 100 bp of each other in G_WOLF coordinates. Any merged loci that contained more than one variant were removed from further processing. An UpSet plot was then created to summarize the pattern of sharing across the sample set using the UpSetPlot module (version 0.9.0) (92).

To estimate a phylogenetic tree from dimorphic SINECs, we encoded presence/absence data for each sample as a sequence in fasta format with a single nucleotide representing a single SINEC locus. We supplemented this with an additional sequence encoding the empty site for each locus which represents the ancestral state. A neighbor joining phylogenetic tree was then estimated from the resulting sequencing using MEGA (v11.0.13) with the p-distance mutation model, which is simply the proportion of sites that differ (93, 94). Uncertainty in tree typology was assessed using 1,000 bootstrap replications (95). The bootstrap consensus tree was rooted using the ancestral empty-site sequence described above. Individual SINEC loci were assigned to specific branches on the inferred tree based on their presence/absence pattern across samples. A second phylogenetic tree was also generated using the above methods on dimorphic LINE-1 insertions. Tables of SINEC and LINE-1 loci aggregated across samples are given in Additional_File2. Final tree visualizations were generated using MEGA and FigTree (93, 96).

### Examination of within-breed variation

To determine whether variation between two GSDs is consistent with that found within other breeds, we performed within-breed comparisons using two GSDs (GSD1 and GSD2), two Labrador Retrievers (LAB1 and LAB2), and two Bernese Mountain Dogs (BMD1 and BMD2). For each pair of samples, the assembly with the smallest segment count was designated as the target, with the other sample being the query.

Segment count was calculated as number of non-‘N’ segments assigned to the autosomes and chrX (Supplemental Data S1). A minimap2 alignment with downstream variant identification was then performed as described above. Within these samples, segmental duplications were identified using BISER and fastCN as described in Nguyen, Blacksmith, and Kidd (80).

### Identification of dimorphic SINEC variants within GSD1

Dimorphic SINECs were identified from a dataset of structural variants identified in GSD1. Specifically, Schall and Kidd identified structural variants in GSD1 by aligning Illumina short-read data and PacBio long-read data generated from GSD1 genomic DNA to the GSD1 assembly (97). Heterozygous autosomal deletions identified in GSD1 were intersected (bedtools intersect -wa -f .9 -r) with SINECs present in GSD1 and absent in G_WOLF (81). These parameters require 90% reciprocal overlap between the dimorphic SINEC and the structural variant.

### Extracting SNVs from inter- and intra-breed comparisons

To identify SNVs relative to G_WOLF or within a single breed, single nucleotide substitutions (i.e., not single base pair deletions or insertions) and regions corresponding to a single aligned segment were extracted from each minimap2 alignment as described above. For each comparison, low confidence regions from the minimap2 target genome were filtered out. The filtered low confidence regions included gaps, BISER (78) and/or fastCN (79) segmental duplications, and loci identified by tandem repeat finder (TRF) (98) in either the query or reference. The low complexity regions were removed from aligned regions with bedtools subtract. Remaining segments were then intersected with SNVs with bedtools intersect to create a final list of SNVs present on the autosomes (81). Note that the autosomal genome segments were only filtered against low complexity regions present in the reference (*i.e.* G_WOLF or the genome with the lower segment count in other comparisons).

### Estimating the rate of canine LINE-1 and SINEC retrotransposition in canines

LINE-1 and SINEC retrotransposition rates were estimated based on average genome divergence times calibrated using SNVs. We combined the filtered number of base pairs in autosomal aligned regions with the number of autosomal SNVs and the estimate canine mutation rate of 4.5x10^-9^ per bp per generation (range 2.6x10^-9^/bp/gen - 7.1x10^-9^/bp/gen) (99) to estimate the number of generations since genome divergence between each sample pair as was done in Nguyen, Blacksmith, and Kidd (80).

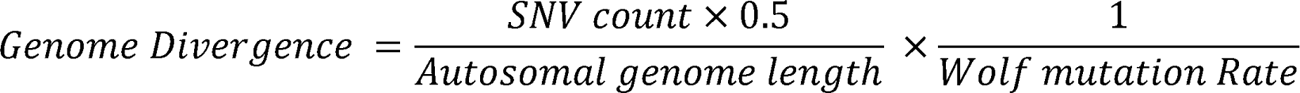

From there, we estimated the rate of LINE-1 and SINEC insertions as the ratio of the number of generations divided by the number of autosomal dimorphic variants.

Estimates were created for dimorphic variants present only in the query, only in target, and the average of both. As an estimate of the uncertainty in the obtained values, we also report insertion rates implied by the range of reported SNV mutation rates. Point estimates of the rate of retrotransposition were also recalculated using only loci which clearly have “high confidence” TSDs of at least 10 bp.

### Identifying LINE-1 3’ transductions

LINE-1s were assessed for the presence of 3’ transductions. First, for each genome, the coordinates of dimorphic LINE-1s were intersected with a bedfile containing the positions of LINE-1s in the genome annotated using bedtools intersect, with option -wao. The longest intersecting annotation was chosen for each locus, and the sequence of candidate 3’ transductions spanning from the end of the LINE-1 annotation to the end of annotated variant, in the same orientation as the LINE-1, were extracted. If the locus possessed a 10 bp or longer TSD, the TSD sequence was removed from the candidate 3’ transduction. Candidates shorter than 25 bp were discarded. The position of repeats and low complexity sequence were identified in each candidate transduction using RepeatMasker (--species dog) and sdust (82, 100). Any candidate that did not have at least 25 bp of unmasked sequence was discarded. The original, unmasked sequence from the remaining candidate 3’ transductions was searched against the G_WOLF reference using blat (version 35, -out=pslx - minIdentity=95) (101). Blat alignments were removed if the match count was less than 25, if the blat target was not on an autosome or chrX, if the target coordinates are within 10 kb of the dimorphic locus, if the span of the aligned target sequence is 100 bp more than or less than the length of the query sequence, if the reported alignment included fewer than 25 bp of unmasked query sequence, if over 50 blat alignments were produced for a single sequence, or if a single locus corresponded to more than five 3’ transductions in a single genome. All blat alignments were aggregated using bedtools sort and bedtools merge (-d 100). The resulting aggregated alignments were parsed to identify if any transduction source corresponds to multiple LINE-1s with transduced sequence in this dataset.

We also identified if blat alignments identified loci that are adjacent to LINE-1s. To do so, all blat alignments mentioned above were aggregated, sorted with bedtools sort, and intersected with G_WOLF LINE-1s using bedtools window (-w 50).

Intersections were then filtered to remove LINE-1s present only in the transduced sequence as detected by blat or LINE-1s which encompass the entire transduced sequence. Remaining alignments were then assessed to note if: (***1***) the transduction was “downstream” of the LINE-1, and (***2***) the orientation of the transduced sequence is in the expected orientation. If both criteria are present, the G_WOLF LINE-1 is considered “adjacent” to the transduction.

Finally, we investigated if any transduced sequences arise adjacent to multiple independent dimorphic LINE-1 insertions. The locations of aligned transductions in G_WOLF coordinates were aggregated, sorted with bedtools sort and merged with bedtools merge (-d 100).

### Rescaling human retrotransposition rates

Both phylogenetic and population modeling estimates of the rate of retrotransposition are highly sensitive to estimates of the rate of SNV mutagenesis. This complicates comparisons as estimates of the human SNV mutation rate have changed over time. Early estimates such as those provided by Nachman and Crowell were around 2.5x10^-8^ per nucleotide per generation (102). However, more recent estimates cluster between 1.1x10^-8^ and 1.45x10^-8^ per nucleotide per generation (103–106). As a result, we rescaled existing rates using a mutation rate of 1.3x10^-8^ mutations per nucleotide per generation.

Additionally, estimates of effective population size (N_e_) are inversely correlated to estimates of the SNV mutation rate. Mallick et al. estimated the rate of human heterozygosity on a population-by-population basis. Heterozygosity was mostly constrained to the interval between .0005 and .001 (107). This estimate places the long-term human effective population size between ∼10,000 and ∼20,000 (107, 108). For clarity, rescaled estimates also use a human effective population size of 15,000, in the center of the range.

## Results

### Pairwise comparisons reveal thousands of dimorphic SINECs

Dimorphic SINEC insertions were identified in a panel of canine genome assemblies. Each sample assembly was aligned as a query to the G_WOLF reference using minimap2 (77). G_WOLF was chosen as the common target because (*1*) wolves are an outgroup relative to breed dogs and Dingoes (109, 110), (***2***) among North American wolves, Greenland wolves have the lowest level of coyote admixture (111), (***3***) the G_WOLF assembly was created using PacBio HiFi sequencing, which has the lowest sequence error rate of available long-read technologies (112, 113), and (***4***) the G_WOLF assembly is highly contiguous, with assembled chromosomes consisting of only 205 contigs (Table 1). For each pairwise genome alignment, structural variants 50 bp or larger, having at least 70% SINEC content, and present on the autosomes (chr1-chr38) or X chromosome were extracted for further comparisons (Fig. 1, Fig. 2). Due to differences in assembly quality and the expected evolutionary history of the X chromosome, only variants on autosomes are considered when comparisons are made across samples.

**Fig. 1:**
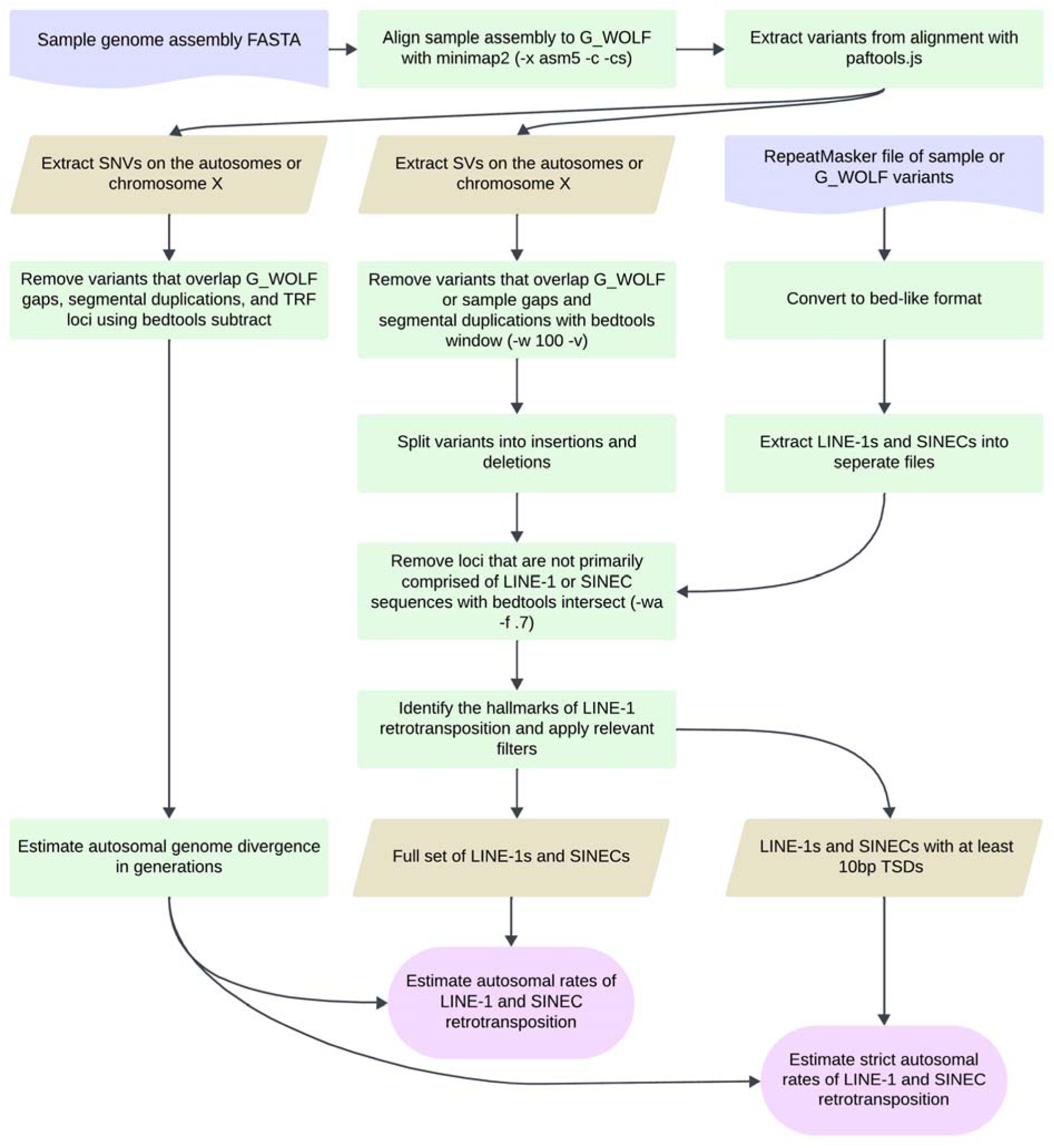
Genome assembly comparison pipeline. The flowchart depicts how single nucleotide and structural variants were identified from canine genome assembly comparisons. Each genome assembly is aligned to the G_WOLF assembly with minimap2 and variants identified from the resulting alignments with paftools.js. SNVs are extracted, filtered, and combined with the fraction of the autosomal genome aligned to estimate the number of generations since genome divergence of the two samples. SVs are extracted, filtered, and split into insertions and deletions and intersected with RepeatMasker annotations to identify candidate LINE-1 and SINEC loci. The SV counts and genome divergence estimates are then compared to calculate an estimated rate of LINE-1 and SINEC insertions per generation. LINE-1 and SINEC containing loci were evaluated for the presence of the hallmarks of retrotransposition allowing for a stricter estimate of the rate of LINE-1 and SINEC retrotransposition. Figure created in Lucid (lucid.co).

**Fig. 2:**
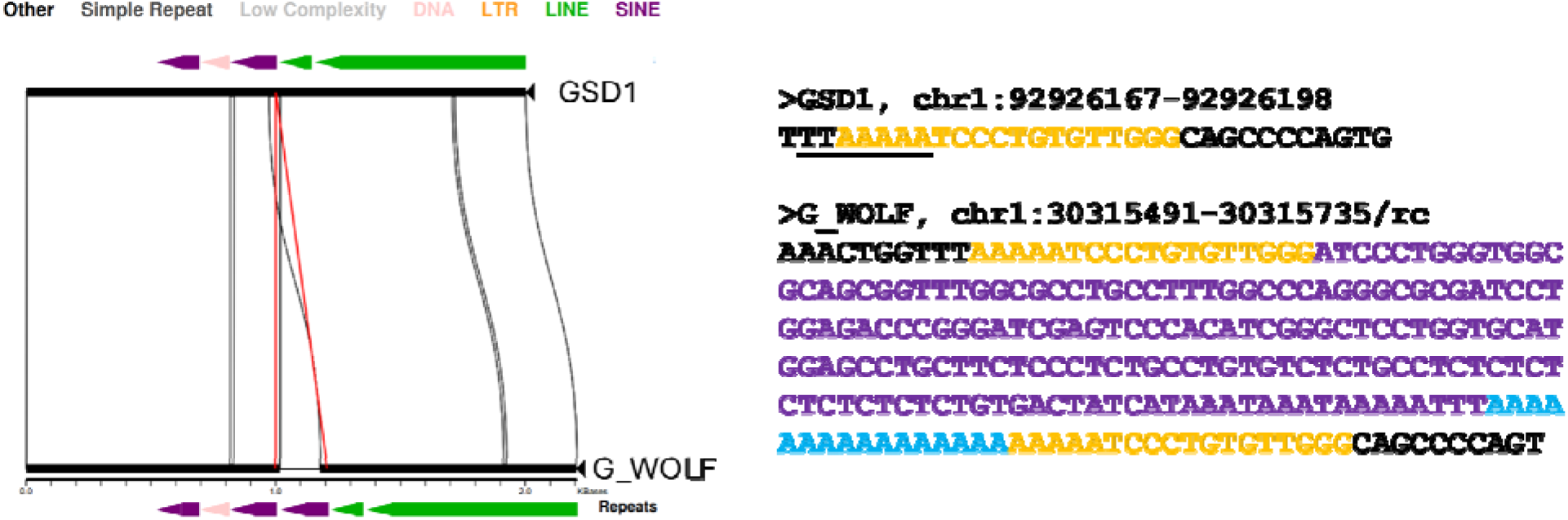
Visualizing a dimorphic SINEC locus. (**A**) A miropeats image shows a dimorphic SINEC locus where the empty site is in GSD1 (top) and the filled site is found in G_WOLF (bottom). Segments present in both GSD1 and G_WOLF are represented as solid black rectangles connected by curved lines. The red lines depict the identified breakpoints for a structural variant present in G_WOLF but absent in GSD1. Repetitive sequence identified by RepeatMasker is depicted as colored arrows, corresponding to different repeat types as indicated. The variable sequence corresponds to a SINEC in the minus strand orientation as highlighted by the left-facing purple arrow. (**B**) Sequences corresponding to filled and empty sites at this locus are shown. Target site duplications are shown in orange in both the filled and empty site. SINEC sequence is shown in purple, poly(A) sequence is in blue, and the inferred LINE-1 endonuclease cleavage site is underlined (5’ TTTTT/AA). The underlined sequence corresponds to the reverse-complement (top-strand) of the cleaved sequence.

On average, each genome comparison identified 25,558 dimorphic SINECs (range: 23,284-27,422) on the autosomes (Fig. 3). The number of SINECs present in G_WOLF but absent from the aligned sample is more consistent across comparisons (mean=12,789, standard deviation (std) 337) than the number of SINECs present in each sample but absent from G_WOLF (mean=12,769, std 1,592). This difference is driven in part by the deficit of dimorphic SINECs detected in BOX (Fig. 3A). The decreased count of SINECs in BOX is found across all autosomes (Fig. S1) and is consistent with a previous description of biased representation of heterozygous SINECs in this assembly (73).

**Fig. 3:**
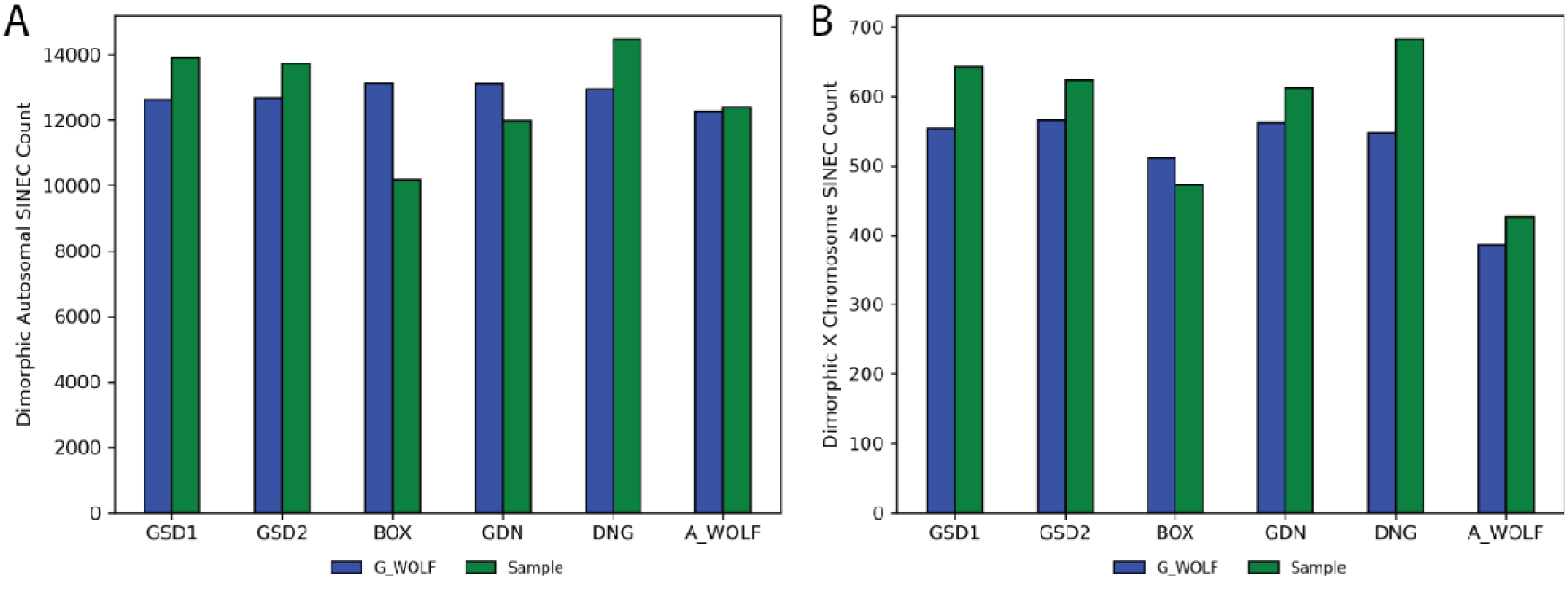
Inter-genome alignments reveal over 20,000 dimorphic SINECs per sample. The bar charts depict the number of dimorphic SINECs identified in each assembly comparison located on the autosomes (**A**) and on chrX (**B**). In each chart, bars represent variants present in G_WOLF, while green bars represent variants present in the sample.

We next refined the coordinates of candidate dimorphic SINECs using the alignment with gap excision (AGE) software (85). After refining the variant coordinates, we searched for the hallmarks of retrotransposition at each locus, which includes the presence of TSDs, 3’ poly(A) tails, and a LINE-1 endonuclease cleavage site (Fig. 4). We began by examining variants present in an alignment between GSD1 and G_WOLF. Dimorphic SINEC loci present in either GSD1 or G_WOLF have similar TSD and 3’ poly(A) lengths. Additionally, loci that possess a ‘high confidence’ (i.e., 10 bp or longer TSDs) contain a sequence profile consistent with the known LINE-1 endonuclease consensus cleavage site (29, 43, 44). Overall, of the variants identified in GSD1, 52.5% have high confidence TSDs and poly(A) tails, 31.2% have only a high confidence TSD, 5.8% have only a high confidence poly(A), 10.5% lack both hallmarks, and 0.1% have ambiguous RepeatMasker orientations (Supplemental Data S2). A comparison of TSD and poly(A) tail lengths indicates that loci with a high confidence TSD possess a wide range of poly(A) tail lengths while loci with short or non-existent TSDs often have short or non-existent poly(A) tails (Fig. S2). Hallmarks are found in similar proportions across each of the analyzed assemblies (Fig. 5). RepeatMasker analysis of the extracted variant sequence shows that 95.1% of dimorphic SINECs present in GSD1 but absent in G_WOLF belong to the SINEC_Cf/2A1 family. Restricting analysis to loci with high confidence TSDs increases this proportion to 96.2%. A similar pattern is found for loci present in G_WOLF but absent in GSD1 (Fig. S3).

**Fig. 4:**
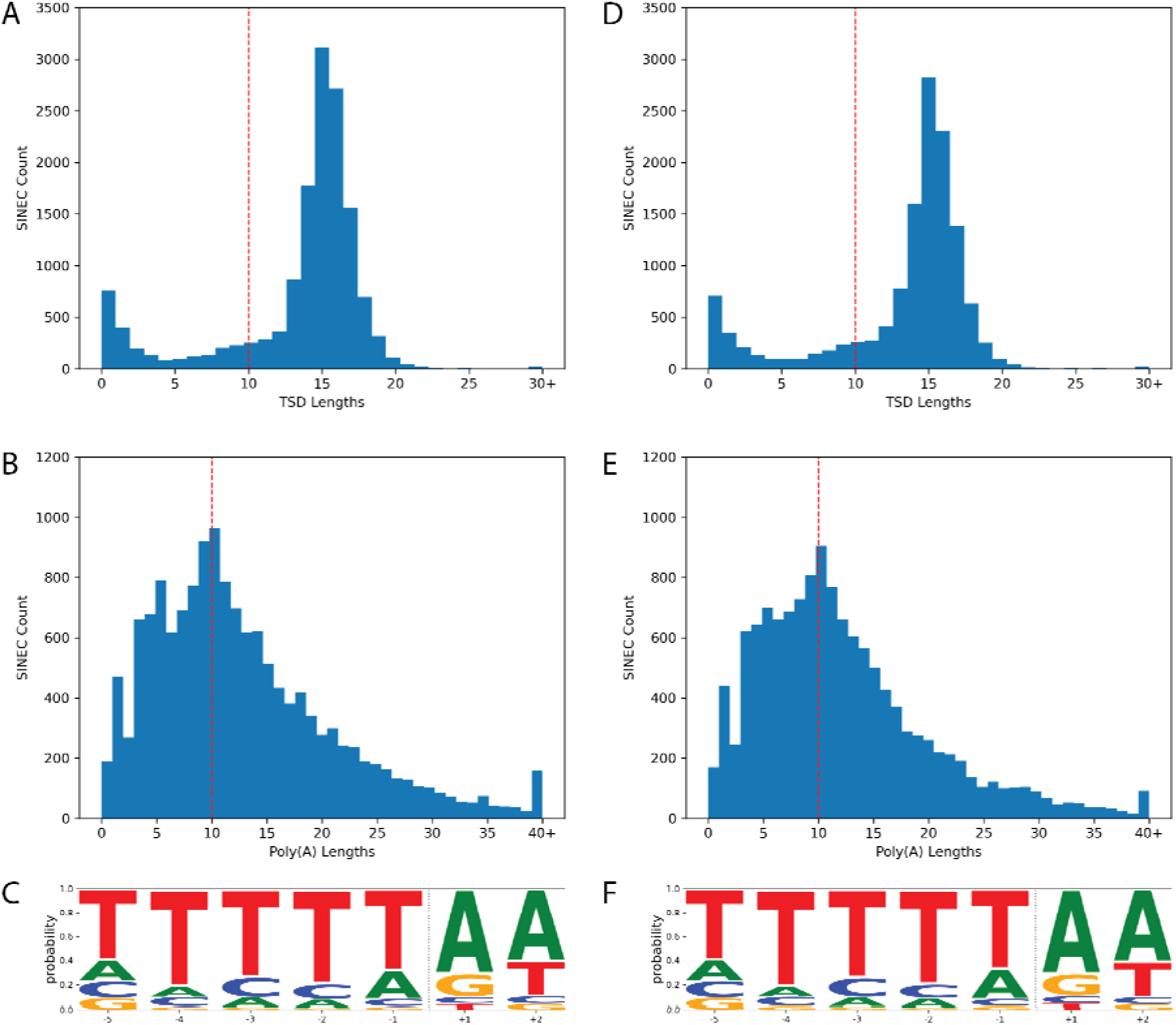
Dimorphic SINECs possess the hallmarks of retrotransposition. The hallmarks of retrotransposition were investigated in SINECs present in GSD1 but not G_WOLF (panels **A-C**) and G_WOLF but not GSD1 (panels **D-F**) according to the GSD1-G_Wolf alignment. Histograms depicting the lengths of identified TSDs and poly(A) tails reveal that most SINECs possess predicted structural hallmarks. A red line depicts the cutoff of 10 bp for high confidence TSDs and 3’ poly(A) tracts (panels **A, B, D, and E**). Logo plots depict that the loci which possess a TSD of at least 10 bp in length tend to possess the canonical LINE-1 EN cleavage site. The x-axis represents the position within the motif, and the dotted vertical line represents the estimated cut site (panels **C and F**). Results are shown for variants on the autosomes and chrX.

**Fig. 5:**
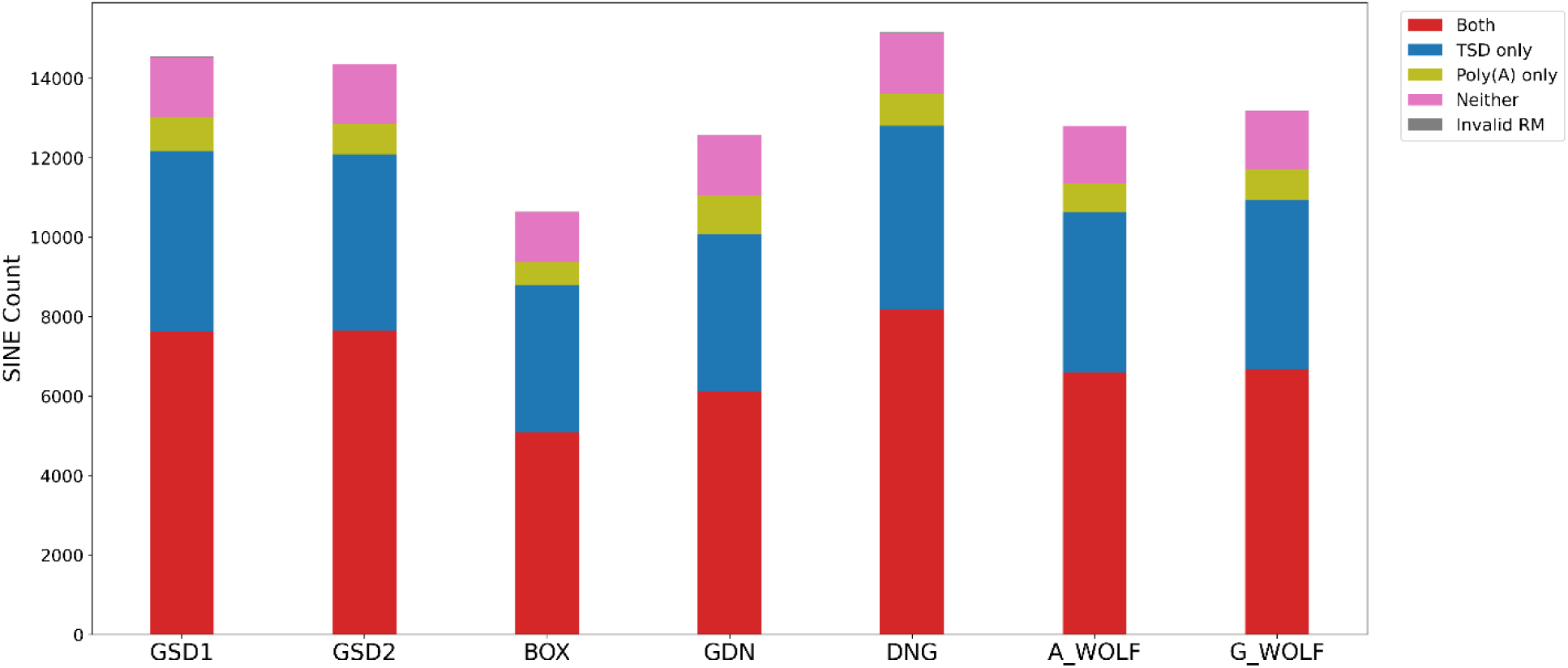
Dimorphic SINECs possess the hallmarks of retrotransposition across samples. All SINEs were categorized as possessing TSDs (>= 10 bp), poly(A) tails (>= 10 bp), both hallmarks, neither hallmark, or having ambiguous RepeatMasker annotations. Stacked bar plots show the distribution of categories across the dataset. The first 6 bars depict variants present in the indicated sample but absent in G_WOLF; the final bar depicts variants present in G_WOLF but absent in GSD1. Loci were only considered “Both” if the poly(A) was not separated by more than 5 bp from the TSD. Results are shown for variants on the autosomes or chrX. If a variant possesses RepeatMasker identified segments in multiple orientations, the variant is placed in the “invalid RM” category.

### High SINEC diversity in canines

To identify variant sharing across the dataset, we identified 2,048,395,076 autosomal bp (in G_WOLF coordinates) that passed callability criteria in each comparison. Of the 51,572 autosomal dimorphic SINEC insertions retained for analysis, 23,865 (46.3%) were present in only one of the seven analyzed assemblies. G_WOLF, A_WOLF, and DNG all possessed more than 5,000 singleton variants. By contrast, each breed dog had between 1,291 and 2,293 singletons, likely a result of the loss of variation during breed formation. BOX had the fewest singletons, and both GSD1 and GSD2 contained fewer singletons than GDN. GSD1 and GSD2 had the most pairwise SINEC variants in common, 1,819, that were exclusively shared, yet each also contained 1,827 and 1,763 singletons. Close behind with 1,794 variants exclusively shared is A_WOLF and G_WOLF. These data highlight the extensive amount of within-breed SINEC variation present in canines (Fig. S4).

We used dimorphic SINECs to infer a neighbor joining tree across the samples, rooting the tree with an ancestral state sequence represented as the empty site for each locus (Fig. 6). As expected, all breed dogs form a group with the two GSDs forming a pair and the Boxer and Great Dane forming a pair. Outside of breed dogs is the Dingo, which is more related to breed dogs than to wolves, but falls outside of the breed dog grouping. The two wolves form a separate taxonomic pair. The tree has a starlike phylogeny with relatively long terminal branches and comparatively shorter internal branches. All nodes had 100% bootstrap support within this dataset except for the node containing GDN and BOX, which had only 76% bootstrap support (Fig. S5). The genotypes of 58.2% of the loci match the inferred tree. The top non-conforming categories include variants present in all samples except A_WOLF (1359, 2.64%), all samples except G_WOLF (947, 1.84%), and present in A_WOLF and DNG (657, 1.27%). A total of 1,827 and 1,763 SINECs were assigned specifically to the GSD1 and GSD2 branches, respectively.

**Fig 6:**
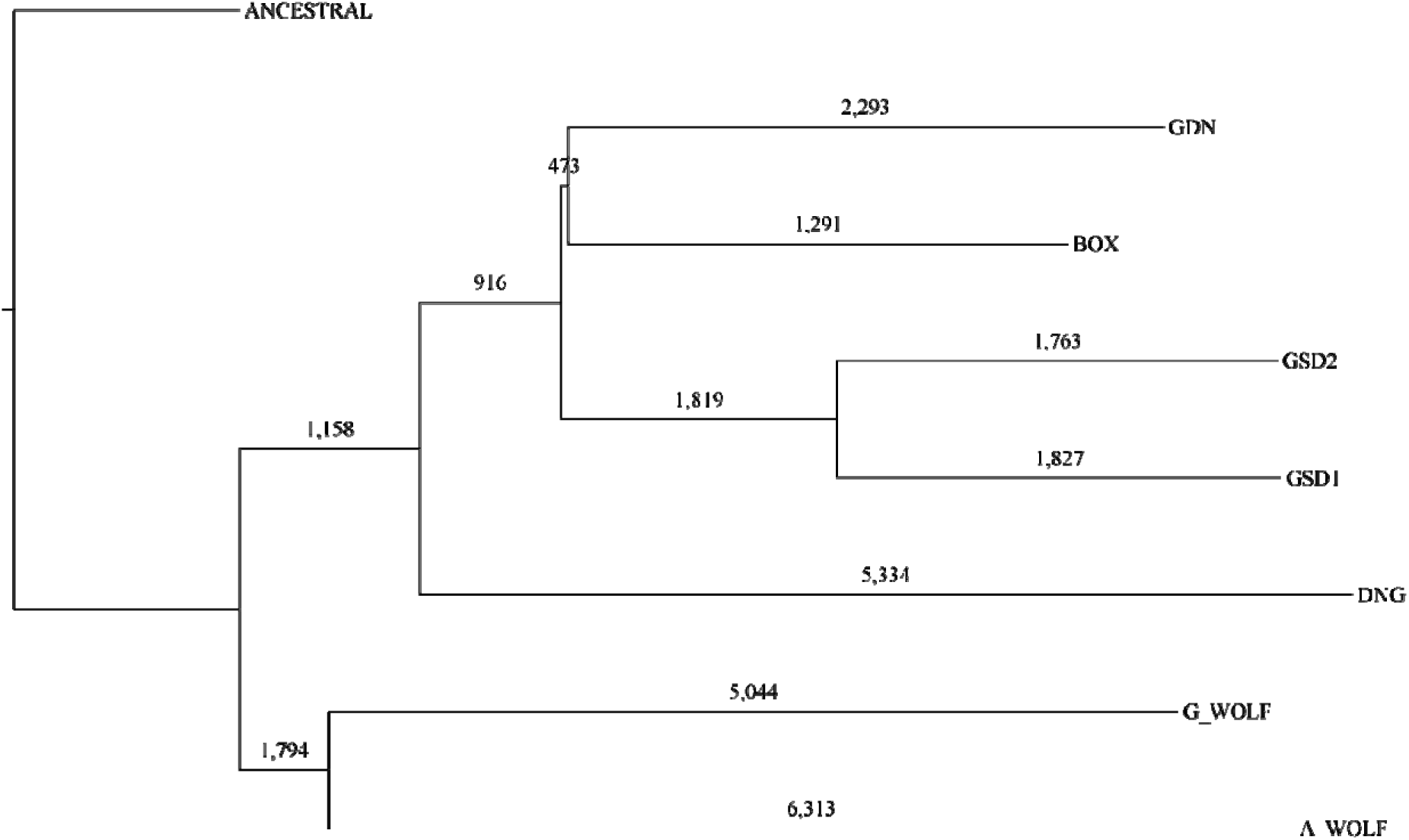
A phylogenetic tree of canine assemblies constructed from dimorphic SINECs. A phylogenetic tree was estimated using a distance matrix created from dimorphic SINEC loci. Trees were rooted on a theoretical ancestral genome for which all dimorphic SINEs are absent. The number of dimorphic SINEC variants is depicted on each branch.

We performed two analyses to confirm the large amount of within-breed SINEC variability. First, we generated within-breed alignments of genome assemblies from the two GSDs, as well as two Labrador Retrievers and two Bernese Mountain Dogs, referred to as GSD1 and GSD2, LAB1 and LAB2, and BMD1 and BMD2, respectively. In each of these comparisons we determined the number of SINECs and SNVs that differ between the assemblies. LABs had the most diversity, with each assembly having ∼6,900 SINECs that were absent in the other. BMDs had the lowest count of dimorphic SINECs (∼4,900 per lineage). Because the amount of within-breed genetic diversity differs, we normalized dimorphic SINEC counts by the number of SNVs found between assemblies, revealing a relatively consistent SINEC to SNV ratio of 0.0056-0.0065. (Fig. S6).

We note that in the GSD comparison there appears to be an increase in dimorphic SINECs between GSD1 and GSD2 relative to the phylogenetic analysis above. Several factors contribute to this finding. First, while there are 3,590 SINECs present exclusively in GSD1 or GSD2 in our dataset, there is a total of 10,569 SINECs that are present in either GSD1 or GSD2 but not both. This includes 6,979 loci that do not fit the inferred tree topology and are not depicted in Fig. 6 showing that this method recapitulates 94% of variants that are identified in the pairwise comparison of GSD1 and GSD2.

Second, we compared SINEC annotations with previously identified deletions identified from GSD1 short- and long-read data aligned to the GSD1 assembly to identify loci where the assembly contains the insertion allele at a heterozygous locus (97). We identified 13,901 autosomal SINECs that were present in GSD1 and absent in G_WOLF. Of these, 4,166 overlapped with heterozygous deletions identified by short-read data and 4,163 overlapped with heterozygous deletions identified by long-read data including 3,756 heterozygous deletions found by both short-read and long-read data. This comparison suggests that ∼30.0% of SINECs with differential presence between GSD1 and G_WOLF are heterozygous in GSD1. Overall, these measures suggest that there is extensive within-breed, and within-individual, SINEC variation in canines.

### A phylogenetic estimate of the rate of SINEC retrotransposition in canines

We estimated the rate of SINEC insertion in canines calibrated by genome divergence implied by a published per-generation SNV mutation rate estimated in wolves. On average, each genome differs from G_WOLF at 4,190,406 autosomal single nucleotide positions (Fig. 7A). Assuming a mutation rate of 4.5x10^-9^ bp/gen (99), and based on the length of the autosomal genome determined as alignable in each comparison, we estimate that average genome divergence of each sample relative to G_WOLF is 228,568 generations (Fig. 7B). Utilizing SINECs present in each sample and absent in G_WOLF, as well as SINECs present in G_WOLF and absent in GSD1, a SINEC insertion is estimated to occur in 1/18.2 (std = 2.0) births. This rate is somewhat deflated due to the low estimated rate of retrotransposition in BOX (1/22.2) births (Fig. 7C). The rate along the G_WOLF branch implied by each comparison is relatively constant, further suggesting that the BOX genome appears to have a biased representation of SINECs (Fig. S7). When only including SINECs with a TSD of at least 10 bp in length, the average point estimate of the rate of retrotransposition was reduced to 1/21.9 births (Fig. 7D). We note that this estimate is highly sensitive to the assumed SNV mutation rate. Using the lower and upper bounds of the reported SNV mutation rate implies a stringent SINEC insertion rate of 1/13.9 to 1/38.0 births (Supplemental Data S3).

**Fig. 7:**
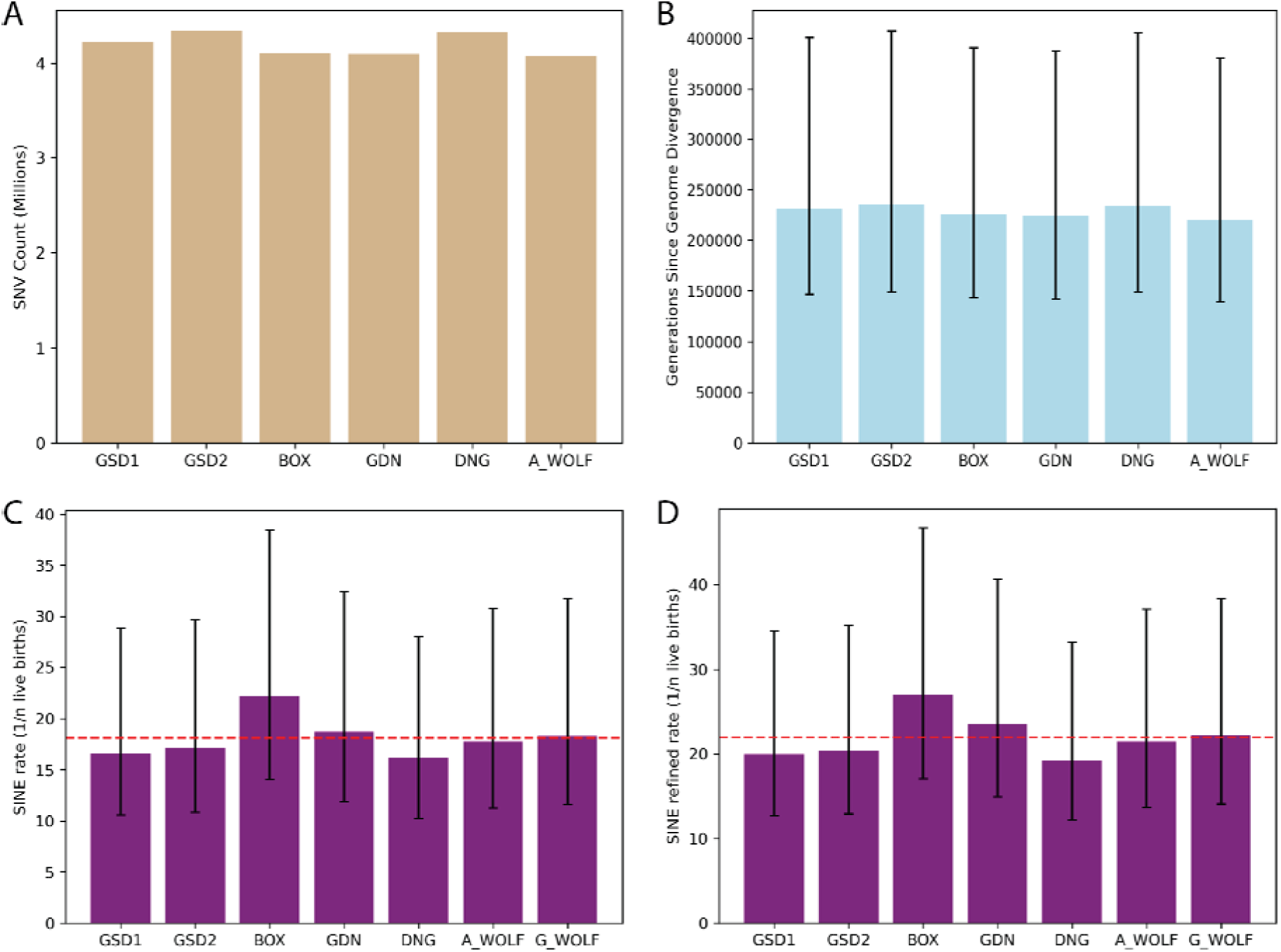
SINECs arise at a rate of ∼1/20 births in canines. Histograms visualize the number of autosomal SNVs between each sample and G_WOLF (panel **A**), as well as the number of generations since divergence using a point estimate of 4.5x10^-9^ bp/generation (panel **B**). The rate of retrotransposition of SINEC was calculated based on all autosomal dimorphic SINEC loci present in each sample and absent in G_WOLF (panel **C**), and a more stringent dataset requiring a TSD of at least 10 bp (panel **D**). For panels C and D, the G_WOLF bar represents variants present in G_WOLF and absent in GSD1. Thus, the corresponding SNVs and generations since divergence can be found in the GSD1 column. A red dotted line shows the average rate of retrotransposition across the dataset. Error bars depict divergence and SINEC retrotransposition rates calculated based on the reported SNP mutation rate confidence interval of 2.6x10^-9^/bp/generation to 7.1x10^-9^/bp/generation.

### Canine samples possess thousands of dimorphic LINE-1 insertions

Using methods similar to those used for SINEC identification, we identified dimorphic LINE-1s among the seven analyzed assemblies. Because deletion of sequence within existing LINE-1 elements could be erroneously identified as a dimorphic element, we implemented two additional criteria to enrich for dimorphic LINE-1s. First, we ran RepeatMasker directly on the inserted sequence and discarded loci that had less than 70% LINE-1 sequence. Second, we removed loci that are embedded within a LINE-1 annotation that do not have a TSD of 10 bp or longer. Using these criteria, an average of 3,497 (range 3,070-3,757) dimorphic autosomal LINE-1 insertions were found in the samples (Fig. S8). In contrast to the profile observed for SINECs, there is no reduction of LINE-1s found in BOX; however, there is a reduction of dimorphic LINE-1 variants found in the A_WOLF assembly. When looking at the variants for which the sample is the filled site, an average of 1.90-fold excess of LINE-1 insertions on the X chromosome relative to the similarly sized chromosome 1 was identified (Fig. S8), consistent with an X-chromosome bias previously observed for LINE-1 (114).

Putative LINE-1 insertions were queried for the presence of the hallmarks of LINE-1 retrotransposition. While the distributions for TSD lengths and endonuclease cleavage site sequences were similar to those discovered in SINECs, a greater proportion of LINE-1 loci have poly(A) tails longer than 40 bp (Fig. S9, Fig. S10). Across the dataset, an average of 43.8% of loci possess both high confidence hallmarks in proximity to each other. Another 27.2% have TSDs only, 10.4% have 3’ poly(A)s only, 13.8% lack both hallmarks, and 4.8% have multiple LINE-1 segments in different orientations for which we did not assess the presence of hallmarks in this analysis (Fig. 8, Supplemental Data S4). An average of 4.1% loci per sample contained two segments annotated as belonging to the same LINE-1 family present in an inverted orientation consistent with twin-priming (47). These potential twin-priming loci were included in the data set but were not processed for the presence of further hallmarks. In GSD1, RepeatMasker analysis of extracted sequence shows that 70.2% of loci belong to the most recent LINE-1 subfamily, L1_Cf, while 21.4% and 4.9% of loci are classified as belonging to the L1MEc and L1_Canis1 subfamilies. The remaining loci were spread across numerous subfamilies with no single subfamily comprising more than 1% of the dataset. Any locus which had segments annotated as belonging to multiple subfamilies by RepeatMasker was reported as unresolved (Fig. S11). When restricting loci to variants which have a high confidence TSD (Fig. 8), the fraction of L1_Cf variants drops slightly to 65.4% while the L1Mec and L1_Canis1 families increase to 25.9% and 5.7%, respectively.

**Fig. 8:**
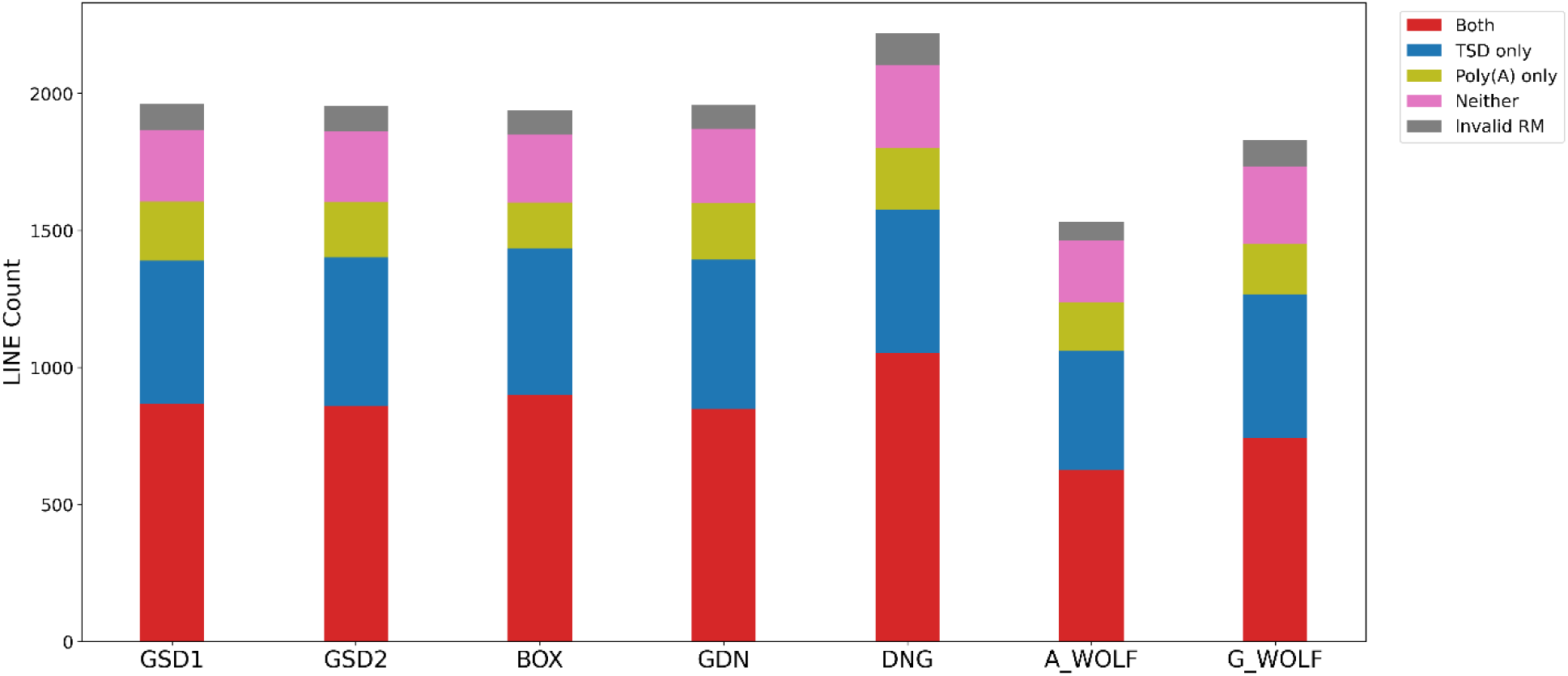
Forty percent of dimorphic LINE-1 loci possess a high confidence TSD and 3’ poly(A) tail. As with SINEs, LINE-1s were categorized as possessing TSDs (>= 10bp), poly(A) tails (>= 10 bp), both hallmarks, neither hallmark, or having ambiguous RepeatMasker orientations. Stacked bar plots show the distribution of categories across the dataset. The first six bars depict variants present in the indicated sample but absent in G_WOLF; the final bar depicts variants present in G_WOLF but absent in GSD1. Loci were only considered “Both” if the poly(A) was not separated by more than 5 bp from the TSD. Results are shown for variants on the autosomes or chrX. If a variant possesses RepeatMasker identified segments in multiple orientations, the variant is placed in the “invalid RM” category.

### Dimorphic LINE-1 insertions possess 3’ transductions

Next, we identified 3’ transductions that could be mapped to the G_WOLF genome from dimorphic loci. When looking at variants present in any sample but not G_WOLF, or G_WOLF but not GSD1 (to avoid counting G_WOLF multiple times in averages), the average genome possessed 63 dimorphic LINE-1 insertions with eligible 3’ transductions that meet our filtering requirements. Sequences were then searched against the G_WOLF reference genome to identify other genomic locations possessing the same sequence (blat hits with >=95% identity). This conservative method identified an average of 35.6 dimorphic LINE-1s per sample with qualifying transductions from the average of 1,913 LINE-1 insertions per genome (Supplemental Data S5).

Focusing on variants present in GSD1 and absent in G_WOLF, a total of 27 dimorphic LINE-1s possessed 3’ transductions which map to G_WOLF. Each of these loci correspond to a single location in the G_WOLF genome. It was noted in Halo et al. that most dimorphic LINE-1 3’ transductions are “parentless” (20). Parentless loci occur when the source of the transduced sequence is not adjacent to a LINE-1, because the “parent” locus is dimorphic in the population. Of the 27 transduction-possessing LINE-1s, only three were 50 bp or less downstream of a LINE-1 found in the G_WOLF genome.

We next identified whether any transduction source corresponds to multiple transduction events in our dataset. Ten such loci exist where one G_WOLF source coordinate corresponds to multiple transductions across our dataset. For two pairs of loci, there were multiple G_WOLF coordinates which could potentially exist as the transduction source. Factoring in the ambiguous loci, a total of eight unique transduction sources were identified which contributed to multiple independent transduction events in the dataset. Some sources, such as G_WOLF chr21:46673641-46673765, correspond to multiple insertions in the same genome. The chr21 locus can be found as transduced sequence in LINE-1 insertions present on DNG chr12 and DNG chr28. Others correspond to multiple loci in different genomes, such as G_WOLF coordinates chr7:21713470-21713969 which correspond to transductions present at four separate locations: chr15 in BOX and DNG, chrs 19 and 21 in DNG, and chr36 in A_WOLF. These data suggest that “hot” LINE-1s have contributed to multiple new insertions in recent canine evolution.

### Utilizing dimorphic LINE-1 insertions to recreate canine phylogeny

LINE-1s present in the 2,048,395,076 autosomal bp that were callable in all genomes were categorized based on presence/absence patterns across the dataset. All samples were included in this analysis, as well as an inferred “ancestral” state which possesses an empty site at all dimorphic LINE-1 loci. In total, 7,428 dimorphic LINE-1 loci were retained on the autosomes with 4,034 (54.3%) being present in only a single sample. (Fig. S12, Fig. S13). A neighbor joining tree was inferred using dimorphic LINE-1s and rooted on the “ancestral” state. The tree possessed the same topology as the SINEC-based tree. Again, all nodes had 100% bootstrap support aside from the GDN-BOX node which had 80% bootstrap support. (Fig. S13). Overall, 65.6% of loci have genotype profiles that fit the inferred tree.

### Relaxed criteria identified additional loci harboring the hallmarks of retrotransposition

It may seem unusual that only ∼51% of SINECs and ∼44% of LINE-1s possess both a TSD and a poly(A) tail. We further investigated if loci that do not possess “high confidence” TSDs and/or poly(A) tails have evidence for the presence of lower confidence hallmarks. To do so, we reanalyzed the variants using less stringent parameters. Because TSDs are frequently described as being 7-20 bp on average, we reduced the minimum TSD size to 7 bp (115). We also looked for evidence of a degraded poly(A) tail by identifying 15 bp windows that contain at least 10 As within 30 bp of a TSD, or the end of the variant if no TSD of at least 7 bp is detected (Figure S14). Application of these relaxed criteria has a dramatic effect on hallmark classification. For SINEC variants, 7,629 (52.5%) variants present in GSD1 and absent in G_WOLF possessed both hallmarks when applying our stringent criteria while 12,625 (∼86.8%) loci had evidence of both hallmarks when applying the relaxed criteria described above, with only 1.3% having neither hallmark. Similar results were observed when reanalyzing LINE-1s present in GSD1 and absent in G_WOLF, with the number of loci possessing evidence of both hallmarks increasing from 866 (44.1%) to 1,404 (71.6%) while the number of loci with neither hallmark was reduced from 260 (13.3%) to 49 (2.5%). Considering only poly(A) tails, 94.8% of SINECs and 72.4% of LINE-1s present in GSD1 without a poly(A) tail found by the initial criteria possess a poly(A) region identified by the relaxed criteria, raising the percent of SINEC and LINE-1s in GSD1 with a poly(A) to 97.8% and 87.6%, respectively. The 12.4% of LINE-1s without an annotated poly(A) include 96 loci (4.9% of the total) with multiple annotated LINE-1 segments present in opposite orientations which were not processed for poly(A) presence.

### Estimating the rate of LINE-1 retrotransposition in canines

Using the SNV mutation and divergence estimates discussed in the SINEC section, we estimated the rate of LINE-1 retrotransposition in canines. We used the same average genome divergence as used in the SINEC analysis. (Fig. 7B). We calculated the retrotransposition rate of each sample relative to G_WOLF and of G_WOLF relative to GSD1, to avoid counting G_WOLF multiple times. The average rate of LINE-1 retrotransposition in our dataset was 1/130.2 live births. DNG had the highest rate of retrotransposition with 1/114.9 live births, while there appears to be a reduction of the rate of LINE-1 retrotransposition in A_WOLF relative to the other samples (Fig. 9A, Fig. S15). To generate a retrotransposition rate using only high confidence LINE-1 insertions, a restricted estimate using variants that possess a TSD of 10 bp or longer was produced. Use of the high-confidence set of LINE-1s reduces the estimate of the rate of retrotransposition to 1/183.9 births. As in the SINEC analysis, the upper and lower bounds of the SNV mutation rate were used to calculate upper and lower bounds of the stringent LINE-1 retrotransposition rate as 1/116.5 and 1/318.2 births, respectively (Fig. 9B, Supplemental Data S6).

**Fig. 9:**
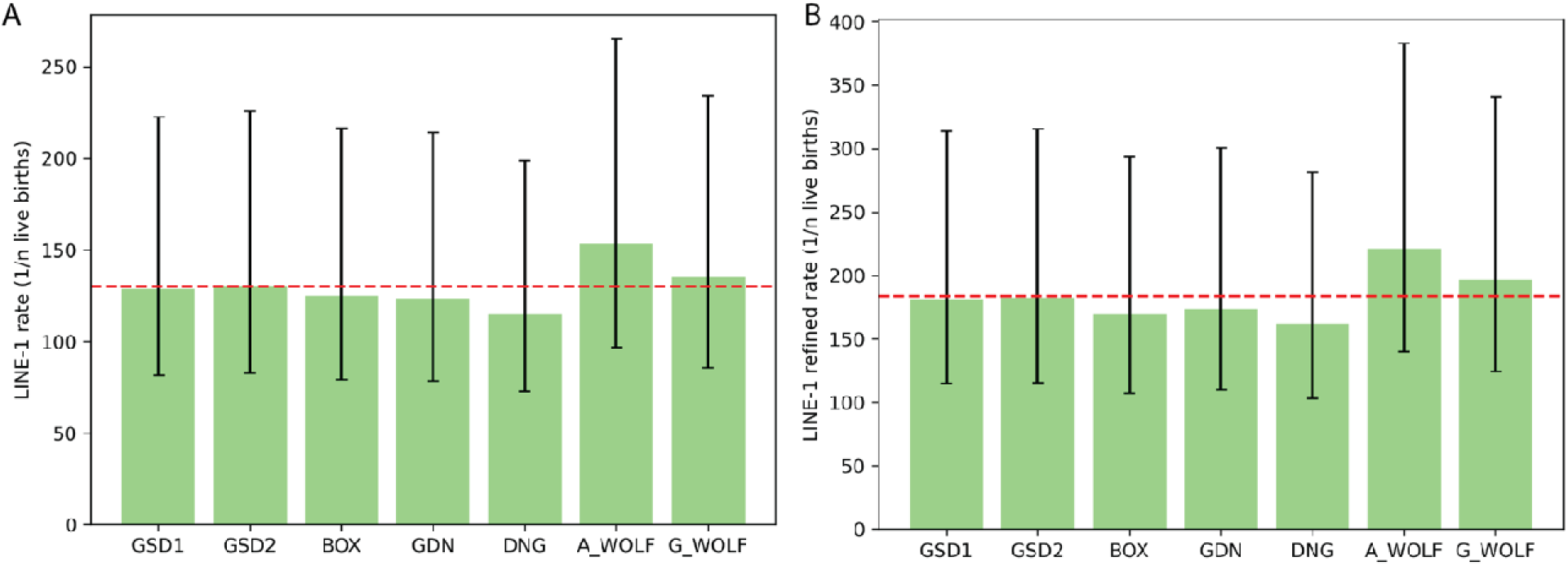
LINE-1s arise at a rate of ∼1/184 births in canines. As in Fig. 7, the rate of retrotransposition of LINE-1 was calculated based on all autosomal dimorphic LINE-1 loci present in each sample and absent in G_WOLF (panel **A**), and a more stringent dataset requiring a TSD of at least 10 bp (panel **B**). G_WOLF bar represents variants present in G_WOLF and absent in GSD1. A red dotted line shows the average rate of retrotransposition across the dataset. Error bars depict divergence and SINEC retrotransposition rates calculated based on the reported SNP mutation rate confidence interval of 2.6x10^-9^/bp/generation to 7.1x10^-9^/bp/generation.

## Discussion

In this study, we compare haploid genome assemblies from four breed dogs, a Dingo, and two wolves to estimate the rate of SINEC and LINE-1 retrotransposition in canines. We estimate a new high confidence LINE-1 and SINEC arising in one out of 184 and 22 live births, respectively. Several factors may bias our estimate. The first is that each reference assembly is a haploid representation of a diploid genome. As a result, these calculations implicitly assume that for heterozygous insertions the filled site and empty site are represented at an equal fraction across the dataset. We note that this assumption is violated in the BOX reference genome, which showed a bias against filled SINEC sites that is not recapitulated for LINE-1 sites (73). The analyzed genomes were generated using a variety of methods for assembling, polishing, gap filling, and scaffolding. BOX was the only reference genome created using the program WTDBG2 and polished with WTDPOA-CNS, which may have influenced the differential inclusion of SINEC variants (116). However, we note that the inferred insertion rate relative to the G_WOLF outgroup is consistent across comparisons, except for SINEC variants in the BOX genome as previously noted. Additionally, the level of within-breed SINEC variation is consistent among German Shepherd Dogs, Labrador Retrievers, and Bernese Mountain Dogs (Fig. S6), a comparison that includes genomes sequenced using Oxford Nanopore Technologies (ONT) long reads. Although these data suggest that biased representation of heterozygous sites is unlikely to majorly skew our findings, future studies that combine phase-resolved diploid assemblies with pangenome analysis approaches may offer a more comprehensive view of canine genome variation.

Although bifurcating trees are not an optimal representation for the complex histories of dogs (117), trees are a useful approach for summarizing the patterns of allele sharing among a collection of samples. The trees we inferred from SINEC and LINE-1 variation recapitulate expected sample relationships with strong bootstrap support. However, only 65.6% of LINE-1 and 58.2% of SINEC loci have presence-absence profiles that match the inferred tree topology, implicating gene flow and/or incomplete lineage sorting of ancestral LINE-1 and SINEC variants as a major contributor to mobile element dimorphism in canines.

Analysis of LINE-1 3’ transductions showed that most transduced sequences that can be aligned to the G_WOLF reference are not located near an existing LINE-1 sequence. This “parentless” configuration suggests the presence of multiple segregating, recently active LINE-1s in canines that are yet to be characterized (20). Tracking of 3’ transductions identified LINE-1s that have given rise to multiple segregating insertions, identifying potentially “hot” canine LINE-1s for future characterization.

We note that the fraction of LINE-1s with 3’ transductions that we identified (∼1.8%) is substantially lower than the ∼15% of full-length LINE-1s with transductions reported in humans (118). There are several reasons for this discrepancy. First, our methodology requires that at least 70% of the inserted sequence corresponds to LINE-1. As a result, truncated LINE-1 insertions that have long 3’ transductions that include non-LINE-1 sequence may fall below the detection threshold. Consistent with this potential bias, we note that transductions appear to be more prominent in longer LINE-1s. For variants present in GSD1 and absent in G_WOLF 1068 (54.4%) and 494 (25.2%) are at least 500 bp and 6000 bp respectively. By contrast, 23 (85.2%) and 12 (44.4%) transduction possessing loci are at least 500 bp and 6000 bp respectively. Second, for a *bona fide* transduction to be identified, the sequence needs to be ≥25 bp and unique (i.e. not composed of low complexity or repetitive sequence). This parameter reduces the number of eligible transductions by ∼40%.

While only an average of 1.8% of loci possessed a mappable transduction in our dataset, this is in line with our previous canine estimates. Our prior comparison between the GDN reference genome and Canfam3.1, which was derived from a boxer, identified 1,121 dimorphic LINE-1s present in one genome or the other. Of them, only 18 (∼1.6%) possessed identifiable 3’ transductions (20). Both analyses yield a much lower proportion of 3’ transductions than identified in human genomes (55, 118). One source of this difference may be LINE-1 3’ UTR sequence divergence between species. We noted several differences between the 3’ UTRs of L1_Cf and L1HS present in RepeatMasker library version dc20170127-rb20170127. Interestingly, the L1HS 3’ UTR possesses a single instance of the “AAUAAA” poly(A) terminator sequence located at the start of the 3’ poly(A) (119). By contrast, the L1_Cf sequence possesses two “AAUAAA” segments located near the 3’ end of the element. The presence of two terminator sequences may contribute to the reduced rate of L1_Cf 3’ transductions.

Although the bulk of our call set contains true insertion variants, some variants that arose through other processes, such as deletions of existing elements, may be included. To ensure that identified variants are indeed LINE-1 mediated, insertions were queried for the hallmarks of LINE-1 retrotransposition. These include the presence of target-site duplications, poly(A) tails, and a LINE-1 endonuclease cleavage site. SINECs overall had a higher fraction of high confidence TSDs than LINE-1s, 83.0% and 71.0%, respectively. There are several reasons why the fraction of variants possessing hallmarks is a lower bound estimate. First, when detecting poly(A) tracts, we required a true homopolymer. Thus, even a single mutation could take a 17 bp homopolymer and split it into two shorter homopolymers; neither of which would qualify as “high confidence”. Similarly, for TSDs, excess mutations can degrade the target site duplication such that the left boundary, right boundary, and empty site are no longer identical. While the AGE aligner does occasionally allow for mismatches among these three sequences, excess differences will prevent the detection of TSDs in this dataset. Additionally, AGE identified a fraction of loci as possessing apparent target site deletions which can arise during TPRT (120, 121), arise from polymorphism at the insertion site, or be assembly errors (Fig. S16). For loci present in GSD1 and absent in G_WOLF, over 80% of loci with no TSD detected by AGE also possess a target site deletion of at least one base. When applying relaxed criteria that includes TSDs of at least 7bp and permits interrupted poly(A) tails, the number of SINEC and LINE-1 variants in GSD1 without either hallmark decreases to ∼1.3% and 2.5%, respectively. The decreased proportion of LINE-1 loci with both hallmarks likely reflects the additional challenges associated with LINE-1 annotation due to the additional variety of structures, including inversion-deletions, found in LINE-1s. Overall, these data suggest that the high stringency filters used for our refined rate estimate are conservative. Other factors may also contribute to the deficit of dimorphic LINE-1s with identified poly(A) tails. For example, we did not systematically process sequences with internal rearrangements. As a result, LINE-1s that integrated via the twin priming mechanism are included in our data set but were not processed for poly(A) tail identification (47).

It is important to note that LINE-1 sequences underwent several additional filters relative to SINECs. This is because a LINE-1 specific class of false positives was identified in our dataset. Specifically, if an existing LINE-1 in a genome undergoes a deletion of internal LINE-1 sequence, the assembly comparison method may incorrectly report a false LINE-1 insertion in other assemblies which do not possess the deletion. *Intra-*LINE-1 deletions are also found in canine SV call sets generated from Illumina data (16), suggesting that this phenomenon reflects real variation and is not an assembly artifact. LINE-1 allelic heterogeneity arising from *intra-*LINE deletions has also been previously reported in humans (122). Thus, we removed any LINE-1 which exists within larger LINE-1 sequences and does not possess a high confidence TSD. While the removal of these variants removed many low-quality loci, remaining variants were not of uniform quality. To reflect this uncertainty, we present estimates for all identified sites as well as only for those sites with an identified TSD.

Unsurprisingly, manual review suggests that some false positives remain in our dataset. For example, a locus found in multiple dogs but absent from G_WOLF (G_WOLF chr6:55577098) likely formed through a mechanism other than retrotransposition. This locus involves sequence from a L1MB4 element, a LINE-1 subfamily that was active ∼150 million years ago (123). The variable sequence is flanked by large segments of matching sequence (130 bp in length) and does not have an inferred cut site that matches the LINE-1 endonuclease consensus sequence. Structural variants such as this locus on chr6 may have formed through homology-mediated processes involving LINE-1-derived sequences and may contribute to the loci with exceptionally long inferred TSDs.

Further analysis shows that simply filtering variants based on the inferred element subfamily is not a robust strategy. RepeatMasker classifies elements based on sequence identity relative to a collection of subfamily consensus sequences and attempts to create a biologically meaningful family assignment from fragmented alignments. For LINE-1s, alignments are processed relative to consensus sequences from different parts of the element (e.g., the 5’ end, the ORF2p region, etc.). The subfamily assignment process may misclassify elements that are fragmented, such as occurs when a mobile element has inserted inside of an existing element (124). Furthermore, RepeatMasker calculates sequence divergence using a set of scoring matrices that vary based on the GC content of the analyzed region. As a result, disparate subfamily assignments may be reported when a sequence is analyzed individually rather than as part of its encompassing chromosomal context. Since the RepeatMasker assignment is based on sequence identity and is not weighted by the presence of specific changes that are diagnostic of subfamily membership, misassignment may occur more frequently when alignments encompass segments with few inter-subfamily differences. Therefore, the subfamily assignments reported in Additional_File1 should be interpreted with caution. A more detailed analysis of canine LINE-1 and SINEC subfamilies, informed by presence-absence patterns found among relevant genome assemblies representing a range of evolutionary distances, may be a fruitful line of future research. Such a detailed subfamily description could then be used to guide the principled assignment of elements into subfamilies, as done by the C*Alu* and LINEu algorithms (125).

In addition to the technical limitations described above, our calculation of the rate of retrotransposition is based on several assumptions. First, the estimate assumes that the rate of LINE-1, SINEC, and SNV mutagenesis has remained constant over the last ∼750,000 years of canine evolution. Violation of this assumption could cause an over- or underestimate of the rate of retrotransposition in canines. Additionally, our phylogenetic estimate assumes that insertions are not subject to genetic selection. Because it is far more likely that new insertions would be deleterious rather than advantageous, the calculated value may underestimate the true rate of *de novo* LINE-1 or SINEC insertion.

The number of generations since divergence is an estimate of the time to genome coalescence between each sample and G_WOLF. This estimate relies upon the number of SNVs identified in each assembly. All assemblies are PacBio derived and the GSD2 and DNG assemblies are supplemented with Oxford Nanopore Technologies (ONT) long reads. Each assembly was also polished using at least one short-read sequencing modality. Each of these differences can affect the base quality throughout the reference assemblies and can propagate further to bias our estimated rates of retrotransposition. It is worth noting, however, that false SNVs would lead to an underestimation of the rate of retrotransposition by artificially increasing the number of generations since divergence and our rate estimates are broadly consistent across samples.

A key parameter in our estimate is the rate of SNV mutation in canines, which we set at 4.5x10^-9^/bp/generation. Using this estimate, the average sample is 229,000 generations diverged from G_WOLF, which corresponds to ∼750,000 years of divergence. We note that this number reflects the average time of lineage coalescence across the genome and, due to the large effective population size found in ancestral wolf populations, is substantially older than the estimated dog-wolf population split time of 10,000-40,000 years ago (126). As described in our previous analysis of retrogenes in canines, this difference is consistent with estimates of wolf demographic history (80). After correcting for differences in the assumed mutation rate, a previous analysis estimated that the ancestors of new-world and old-world wolves had an effective size (Ne) of 143,000 (126). This large size corresponds to an expected time to coalescence of two lineages in the ancestral population of 286,000 generations. After accounting for the additional divergence following population separation, this expectation is broadly consistent with the divergence time we inferred from genome assembly comparisons.

The largest source of error in our estimates of the LINE-1 and SINEC retrotransposition rates is the assumed SNV mutation rate. The value we used was derived from Illumina sequencing of a wolf pedigree with four offspring. The Koch et al. point estimate has a large confidence interval (2.6x10^-9^-7.1x10^-9^) and is broadly consistent with the rate inferred from analysis of an ancient wolf genome (2, 99). Recently, Zhang et al. estimated a SNV mutation rate of 4.89x10^-9^ per bp per generation based on Illumina sequencing of 404 dog trios (127). The authors noted that the dog SNV mutation rate is correlated with paternal age and may also show differences across breeds.

We utilized a phylogenetic approach to estimate the rate of canine LINE-1 and SINEC retrotransposition. This method is based on the total number of insertions that occurred in two samples and is calibrated by a known SNV mutation rate. Other approaches to estimate the rate of retrotransposition have been employed in other species. These include direct estimates obtained through pedigree sequencing and population modeling approaches based on the number of insertions found in a population (i.e., Watterson’s estimator (128)). Population modelling approaches require an estimate of the effective population size of the studied group, a value that can be sensitive to modelling assumptions and is itself often calibrated using a SNV mutation rate. Although conceptually simple, direct pedigree estimates are sensitive to filtering strategies and are influenced by both false positive calls in the offspring and false negative calls in the parents.

To place our estimates of the rate of retrotransposition in canines in a broader context, we compared them to retrotransposition rates estimated in humans (Table 2). We recalibrated published rate estimates to use the smaller human SNV mutation rate implied by pedigree sequencing studies. After recalibration, the estimated rate of *Alu* retrotransposition in humans (∼1/36 – ∼1/40 births) is approximately half of the SINEC rate we estimate for canines. The human LINE-1 insertion rate estimated by phylogenetic or population modelling approaches (∼1/210 - ∼1/360 births) is slightly smaller than our estimate of the rate of LINE-1 insertions in dogs. However, the estimate obtained from human pedigrees (∼1/63) is markedly higher. The extent to which technical factors (i.e., false positives and false negatives) and biological factors (i.e., strong negative selection against new LINE-1 insertions) contribute to differences in the estimated rates is unclear. The modest differences in retrotransposition rates estimated for dogs and humans suggest that the striking levels of canine SINEC and LINE-1 dimorphism reflect long-standing genetic variation that has been segregating in canines throughout their history.

**Table 2:** Recalibrated estimates of LINE-1 and *Alu* retrotransposition in humans.

| Author and Year | Method | Element<br>Type | Published<br>Rate<br>Estimate | Rescaled<br>Rate<br>Estimate |
| --- | --- | --- | --- | --- |
| Xing et al., 2009 | Phylogeny | LINE-1 | 1/212 | 1/360 |
| Feusier et al.,<br>2019 | Pedigree | LINE-1 | 1/63 | 1/63 |
| Ewing &<br>Kazazian, 2010 | Population<br>Modeling | LINE-1 | 1/140 | 1/210 |
| Xing et al., 2009 | Phylogeny | *Alu* | 1/21 | 1/36 |
| Feusier et al.,<br>2019 | Pedigree | *Alu* | 1/40 | 1/40 |

Published rates were obtained directly from the indicated manuscripts. Rescaled rates were calculated based on a human mutation rate of 1.3x10^-8^ mutations per bp per generation and an N_e_ of 15,000.

## Conclusions

By comparing seven canine genome assemblies we identified a total of 7,428 dimorphic LINE-1s and 51,572 dimorphic SINECs present in regions resolved in all genomes. Insertion allele sharing profiles reflect known sample relationships and reveal substantial within-breed variation. Calibrating estimates using a previously estimated single nucleotide mutation rate, we estimate that the rate of SINEC-1 and LINE-1 and retrotransposition is 1/22 and 1/184 births over recent canine evolution. The rate of SINEC insertions in canines is approximately two times larger than the rate estimated for the *Alu* element in humans, while the estimated canine LINE-1 insertion rates are comparable, suggesting that a large pool of segregating insertions have been maintained throughout canine evolution.

## Supporting information

Key to Additional_File1

## Author Contributions

M.S.B., J.V.M., and J.M.K. designed the study and interpreted the results, M.S.B. and A.K.N performed analyses, M.S.B. and J.M.K drafted the manuscript. M.S.B., A.K.N., J.V.M. and J.M.K. edited the manuscript.

## Data Availability

Final outputs from locus processing including variant location, hallmark information, and element subtype are reported in Additional_File1 and Additional_File2 included online with this manuscript. Genome assemblies are available at the accessions given in Supplemental Data S1. Relevant code for this manuscript is available at https://github.com/mblacksmith1996/Canine_LINE1_SINEC_Scripts.

## Acknowledgements

We thank the authors of multiple canine genome assemblies for making their data available to the community. We also thank Emily Koch and Ryan E. Mills for helpful discussion and comments on previous versions of this study and two anonymous reviewers for constructive comments that improved our study.

