## Supplemental Figures for "A phylogenetic estimate of canine retrotransposition rates based on genome assembly comparisons"

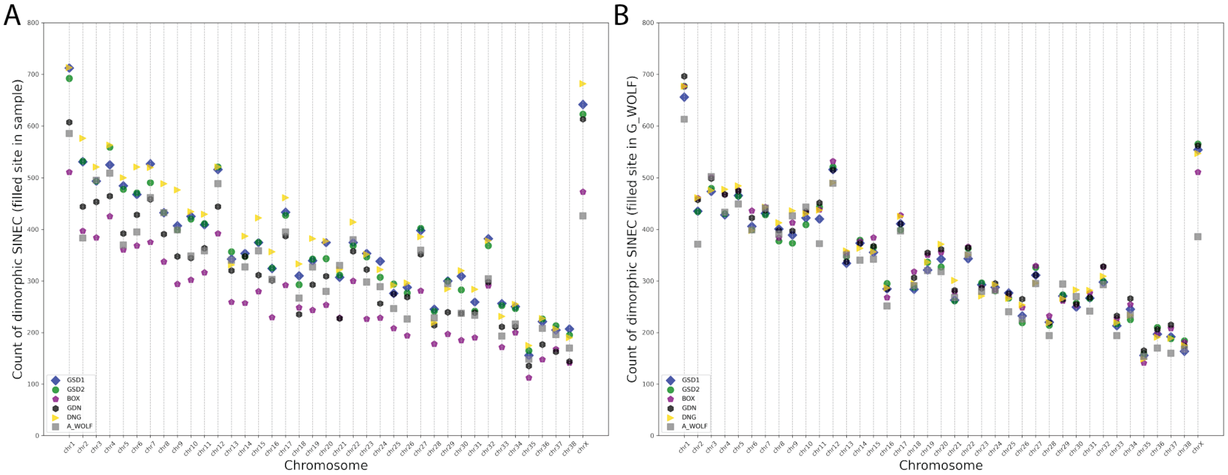


**Fig. S1: SINEC insertions by chromosome.**

Dot plots display the number of SINEC insertions detected per chromosome in each genome comparison. Chromosomes are ordered by autosome number followed by chrX. SINECs present in the sample (panel **A**) and present in G_WOLF (panel **B**) are shown in separate plots.

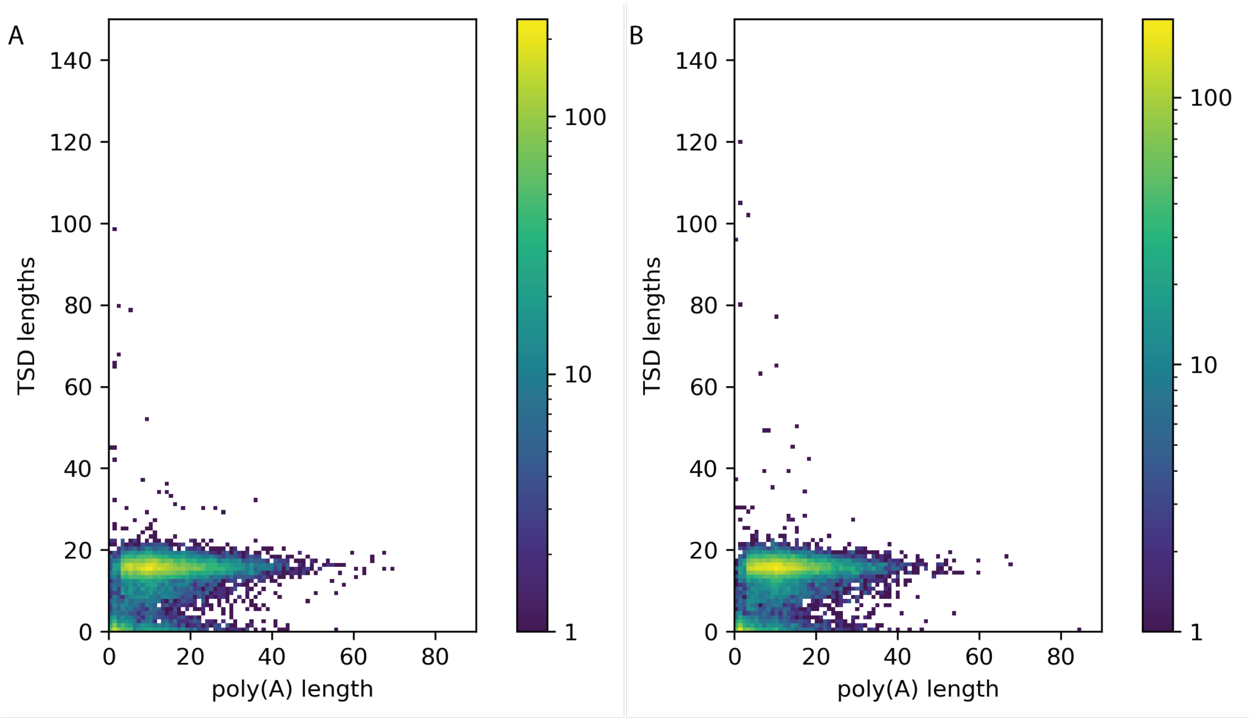


**Fig. S2: Correlating SINEC TSDs and poly(A) tracts.**

A heatmap displays the correlation between the lengths of TSDs and 3' poly(A) sequences for SINECs in GSD1 but not G_WOLF (panel **A**) and G_WOLF but not GSD1 (panel **B**). Each square represents a single bp resolution of TSD and poly(A) lengths. A single G_WOLF variant is not depicted which has a TSD length of 191 bp. Results are shown for variants on the autosomes or chrX. Color bar indicates the count of sites in each coordinate.

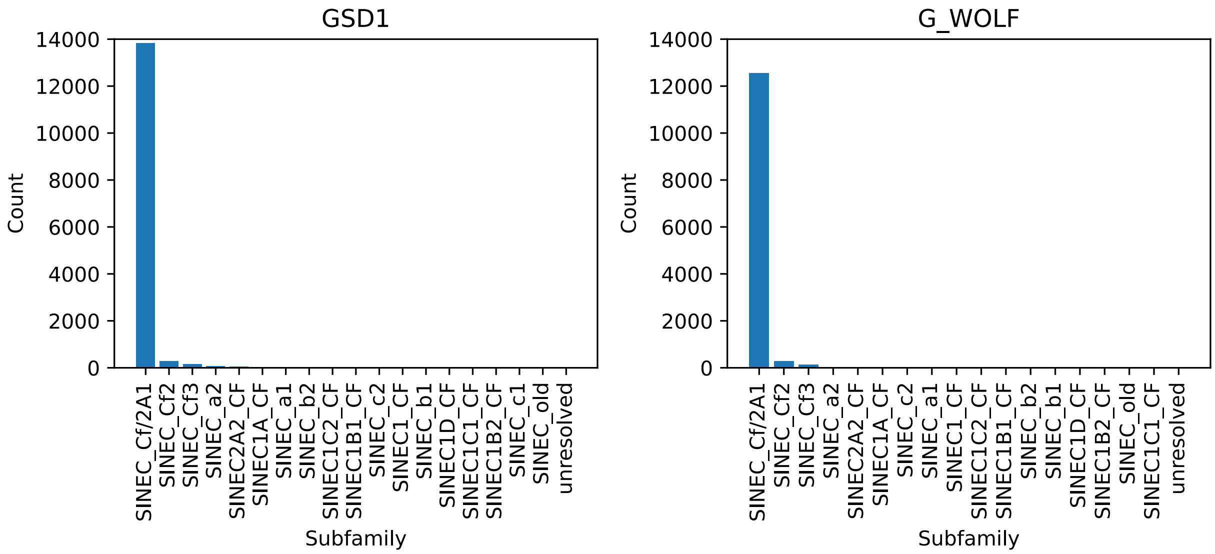


**Fig. S3: Nearly all dimorphic SINECs belong to the SINEC_Cf/2A1 subfamily.**

Histograms of SINEC subfamily as identified by RepeatMasker applied to extracted variant sequences. If multiple different subfamilies are detected within a single locus, the subfamily type is listed as unresolved. SINEC variants present in GSD1 but not G_WOLF (**left**) and G_WOLF but not GSD1 (**right**) are displayed. Results are shown for variants on the autosomes or chrX.

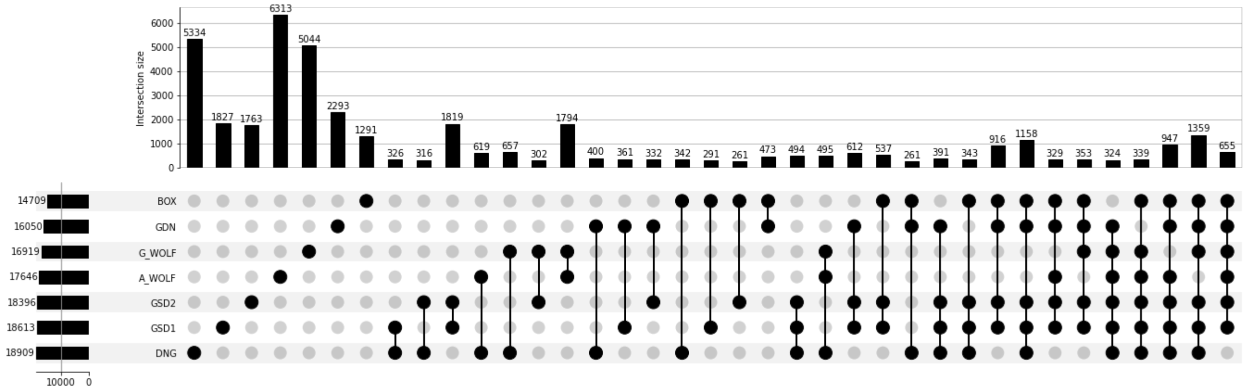


**Fig. S4: UpSet plot of dimorphic autosomal SINEC variants.**

An UpSet plot depicts SINEC variant sharing across samples. Filled dots represent that the variant is present in the indicated sample. Counts are provided for each category above their corresponding bar. Any categories representing less than 0.5% of the dataset are not included in the plot.

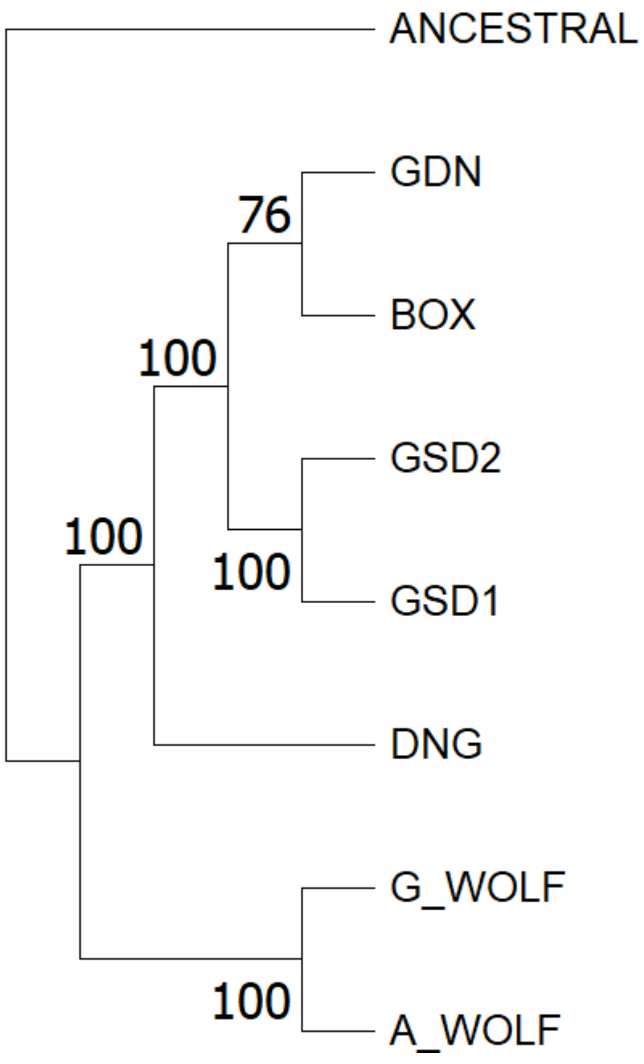


**Fig. S5: Phylogenetic tree developed from dimorphic SINECs is of high confidence.**

Bootstrap support from 1,000 runs is displayed on a phylogenetic tree with the same topology as Fig. 6.

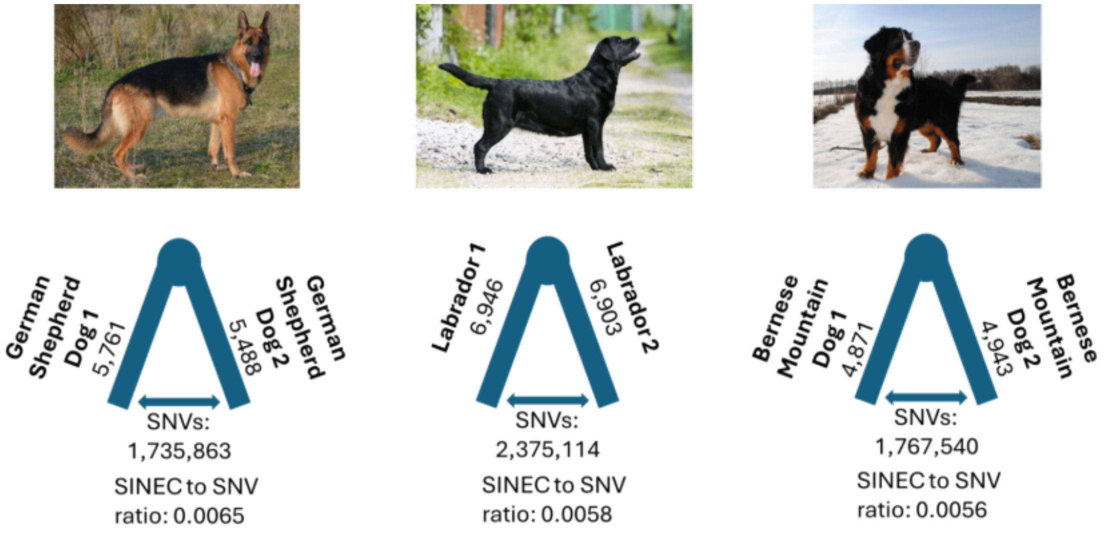


**Fig. S6: Consistent levels of SINEC and single nucleotide diversity are found within three breeds.**

Dimorphic SINECs and SNVs were identified between assemblies of two German Shepherd Dogs (**left**), Labrador retrievers (**center**), and Bernese Mountain dogs (**right**). In each comparison, the number of autosomal dimorphic SINECs, as well as the number of SNVs, are shown. Below each triangle is the SINEC to SNV ratio, which is the total number of dimorphic SINECs divided by the total number of SNVs. Representative images of a dog from each breed obtained from Wikimedia Commons are shown. Original images can be accessed at the following links: https://commons.wikimedia.org/wiki/File:German-shepherd-4040871920._(2).jpg, https://commons.wikimedia.org/wiki/File:Labr.jpg, and https://commons.wikimedia.org/wiki/File:Minelanda_Moon_Russia.JPG.

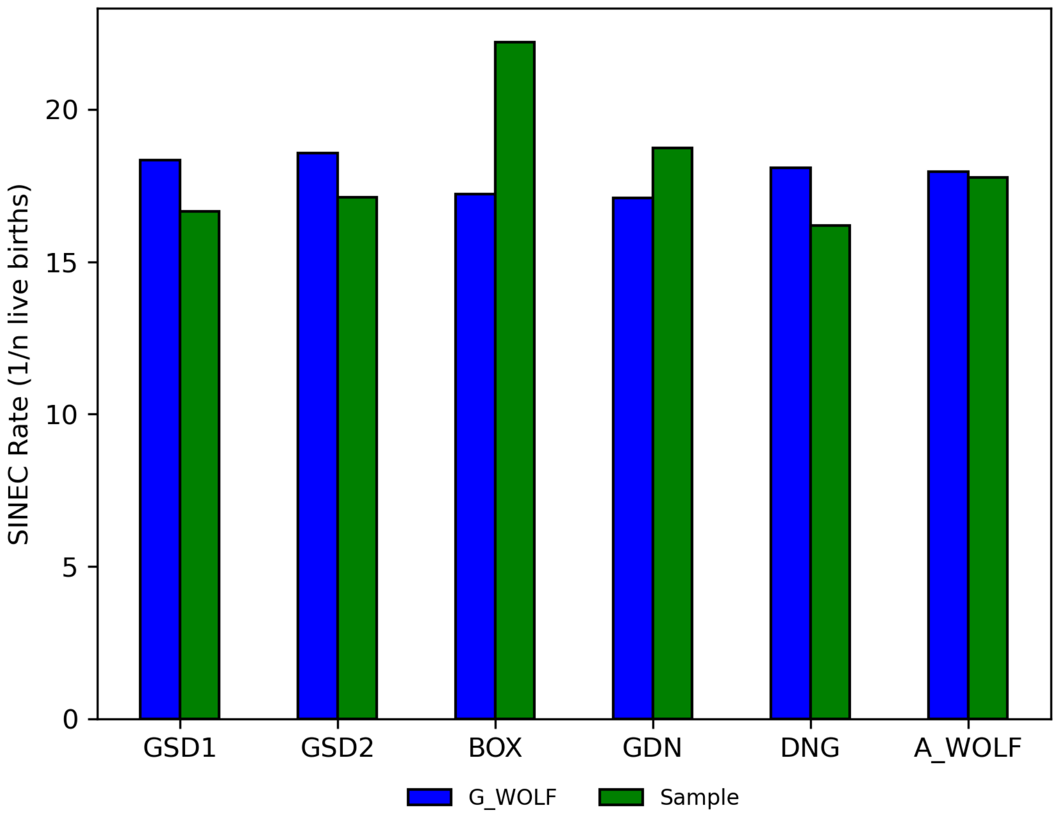


**Fig. S7: SINEC retrotransposition is more variable among queried genomes than G-WOLF.**

A bar plot depicts inferred SINEC insertion rates based on elements that are present in G_WOLF and not the queried sample (**Blue**) and in the sample but not G_WOLF (**Green**). Estimates are based only on autosomal data and assume a SNP mutation rate of $4.5x10^{-9}$/bp/generation.

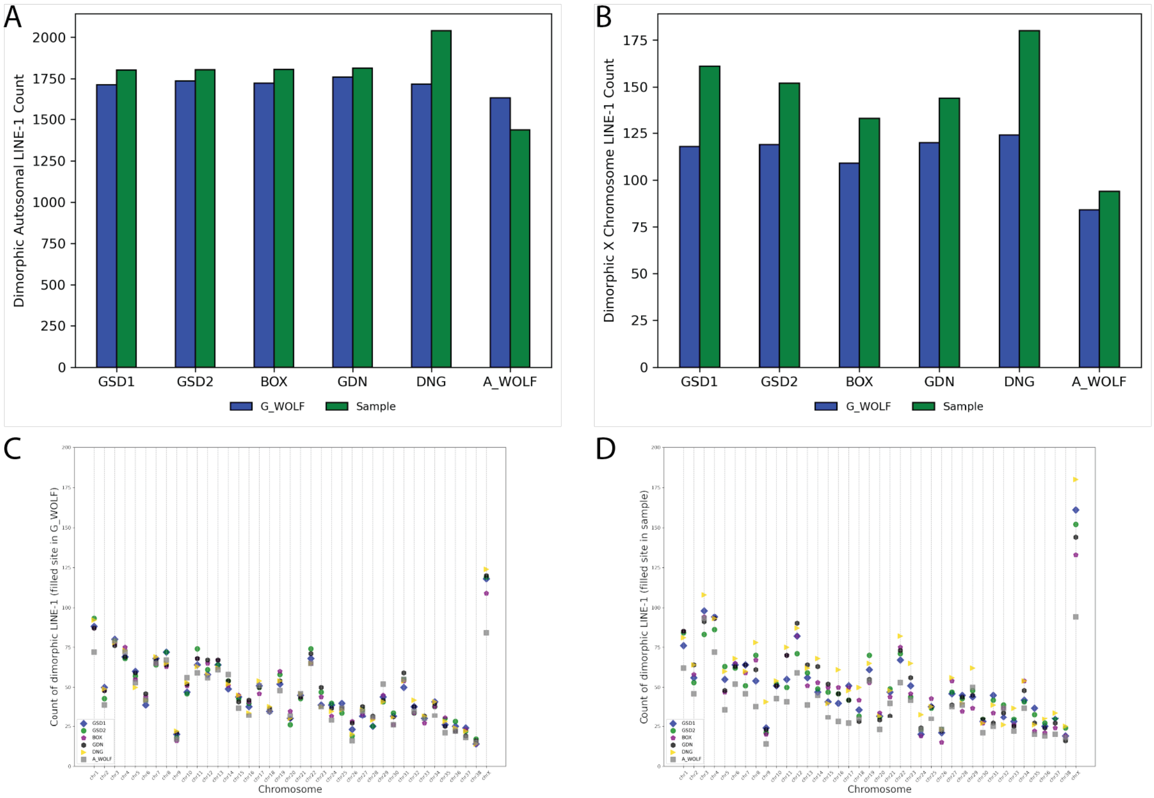


**Fig. S8: Dimorphic LINE-1s are numerous across canine samples.**

Bar charts depict the number of dimorphic LINE-1s identified in each assembly located on the autosomes (panel **A**) and on chrX (panel **B**). In each chart, bars represent variants present in G_WOLF, while green bars represent variants present in the sample. A_WOLF possesses reduced LINE-1 content on the X chromosome. Dot plots improve resolution by showing the number of variants on each autosome and chromosome X for variants in the sample and not G_WOLF (panel **C**) and in G_WOLF but not the sample (panel **D**).

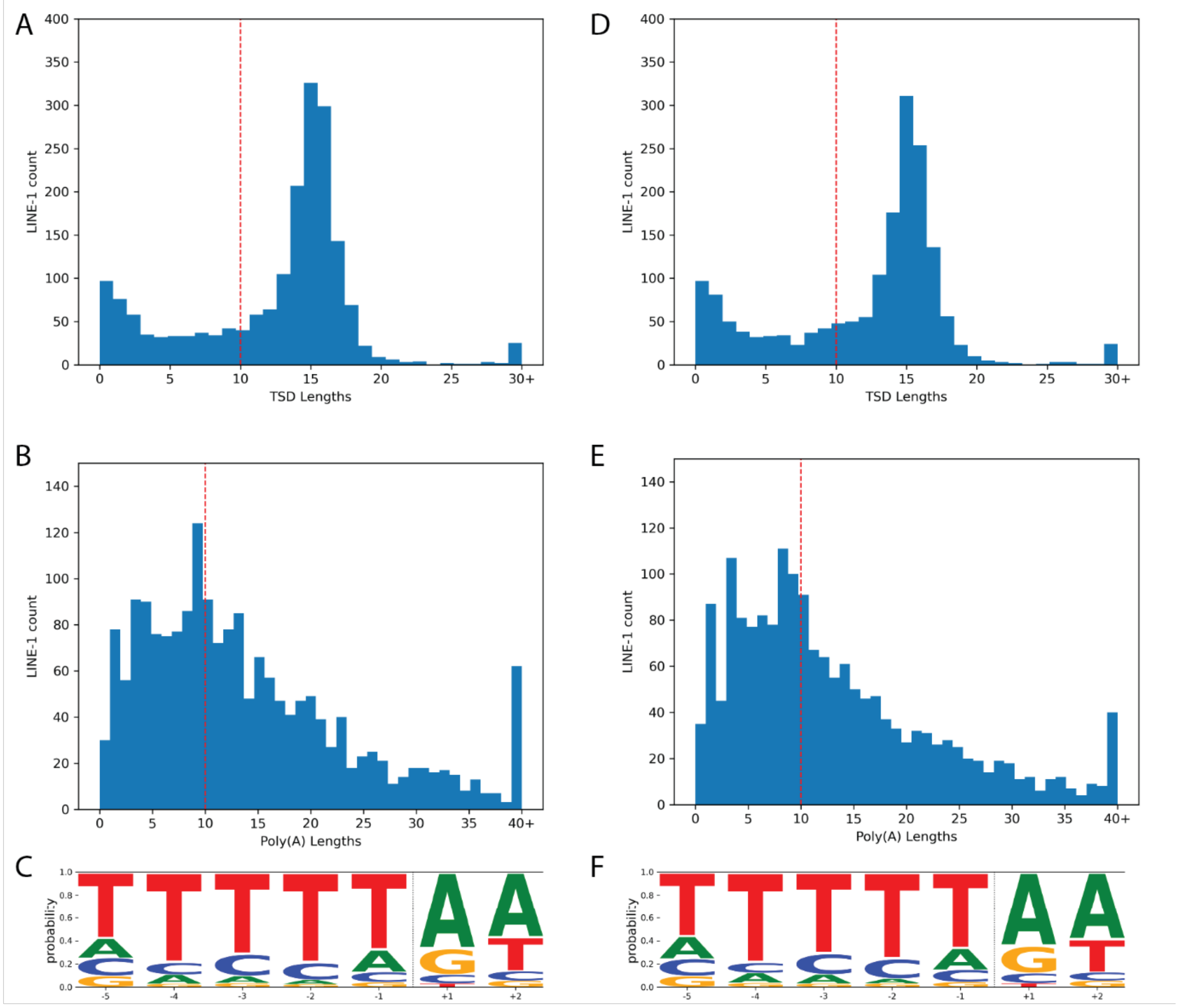

A
LINE-1 count
TSD Lengths
0 5 10 15 20 25 30+
0 50 100 150 200 250 300 350 400
D
LINE-1 count
TSD Lengths
0 5 10 15 20 25 30+
0 50 100 150 200 250 300 350 400
B
LINE-1 count
Poly(A) Lengths
0 5 10 15 20 25 30 35 40+
0 20 40 60 80 100 120 140
E
LINE-1 count
Poly(A) Lengths
0 5 10 15 20 25 30 35 40+
0 20 40 60 80 100 120 140
C
probability
0.0 0.2 0.4 0.6 0.8 1.0
-5 -4 -3 -2 -1 +1 +2
F
probability
0.0 0.2 0.4 0.6 0.8 1.0
-5 -4 -3 -2 -1 +1 +2

**Fig. S9: Most LINE-1 insertions possess high confidence TSDs and/or poly(A) tracts.**

The hallmarks of retrotransposition were investigated in LINE-1s present in GSD1 but not G_WOLF (panels **A-C**) and G_WOLF but not GSD1 (panels **D-F**) according to the GSD1-G_Wolf alignment. Histograms depicting the lengths of identified TSDs and 3' poly(A) tails reveal that most LINE-1s possess hallmarks. A red line depicts the cutoff of 10 bp for high confidence TSDs and poly(A) tracts (panels **A, B, D**, **and E**). Logo plots depict that loci which possess a TSD of at least 10 bp in length possess the canonical LINE-1 EN cleavage site. The x-axis represents the position within the motif, and the dotted vertical line represents the estimated cut site (panels **C and F**). Results are shown for variants on the autosomes and chrX.

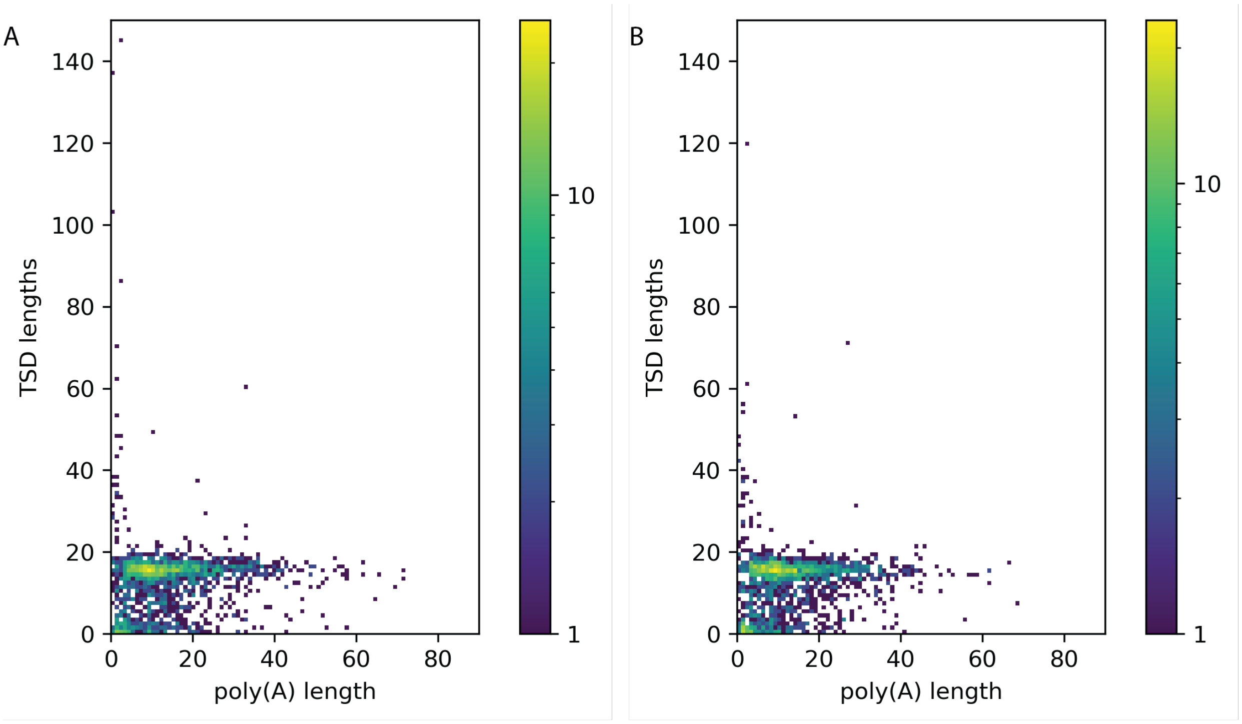


**Fig. S10: The poly(A) tails of dimorphic LINE-1 insertions are more variable than TSDs.**

A heatmap displays the correlation between TSDs and poly(A)s for LINE-1s in GSD1 but not G_WOLF (panel **A**) and G_WOLF but not GSD1 (panel **B**). Each square represents a single bp resolution of TSD and poly(A) lengths. Two outliers are not included in the image: (***1***) A GSD1 variant with a 307 bp TSD; and (***2***) A G_WOLF variant with a 158 bp TSD. Results are shown for variants on the autosomes or chrX. Color bar indicates the count of sites in each coordinate.

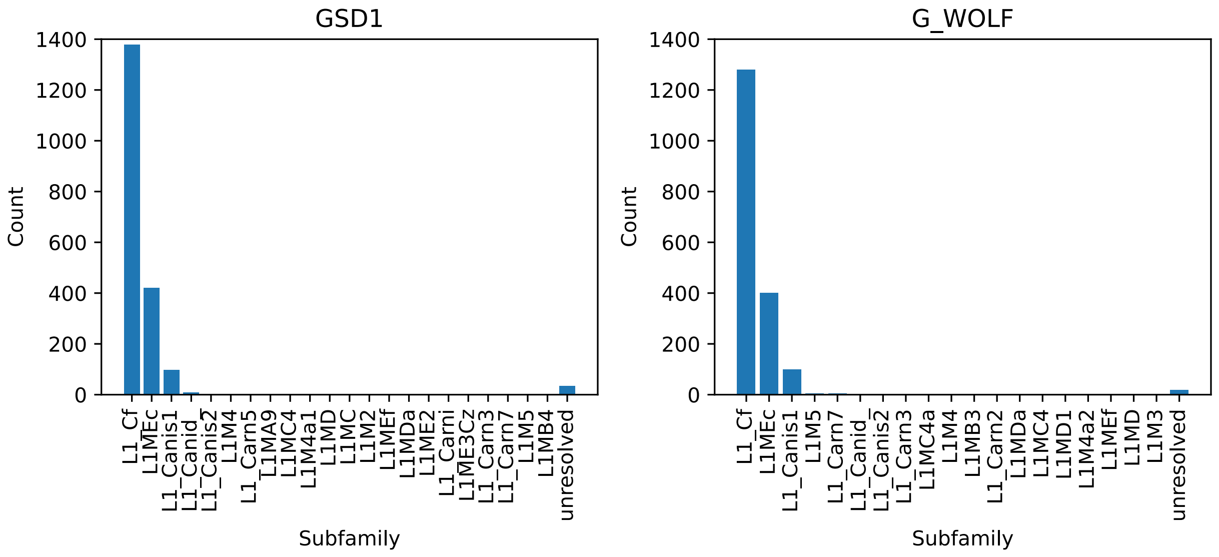


**Fig. S11: The majority of dimorphic LINE-1 insertions in GSD1 and G_WOLF are of the most recent subfamily, L1_Cf.**

Histograms of LINE-1 subfamily as identified by RepeatMasker applied to the extracted sequence of each variant. If multiple different subfamilies are detected within a single locus, the subfamily type is listed as unresolved. LINE-1 variants present in GSD1 but not G_WOLF (**left**) and G_WOLF but not GSD1 (**right**) are displayed. Results are shown for variants on the autosomes or chrX.

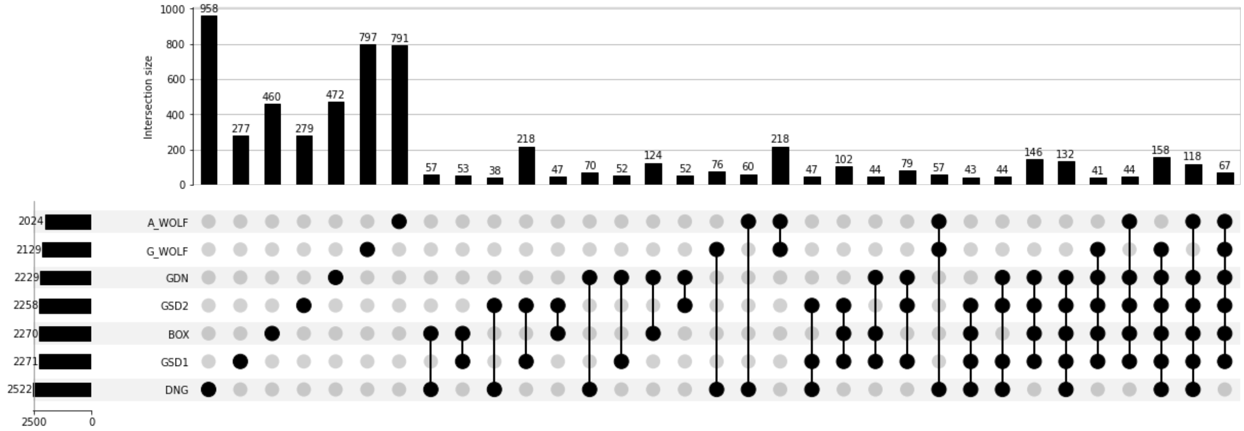


**Fig. S12: Dimorphic LINE-1 sharing across samples.**

An UpSet plot depicts LINE-1 variant sharing across samples. Filled dots represent that the variant is present in the indicated sample. Counts are provided for each category above their corresponding bar. Any categories representing less than 0.5% of the dataset are not included in the plot.

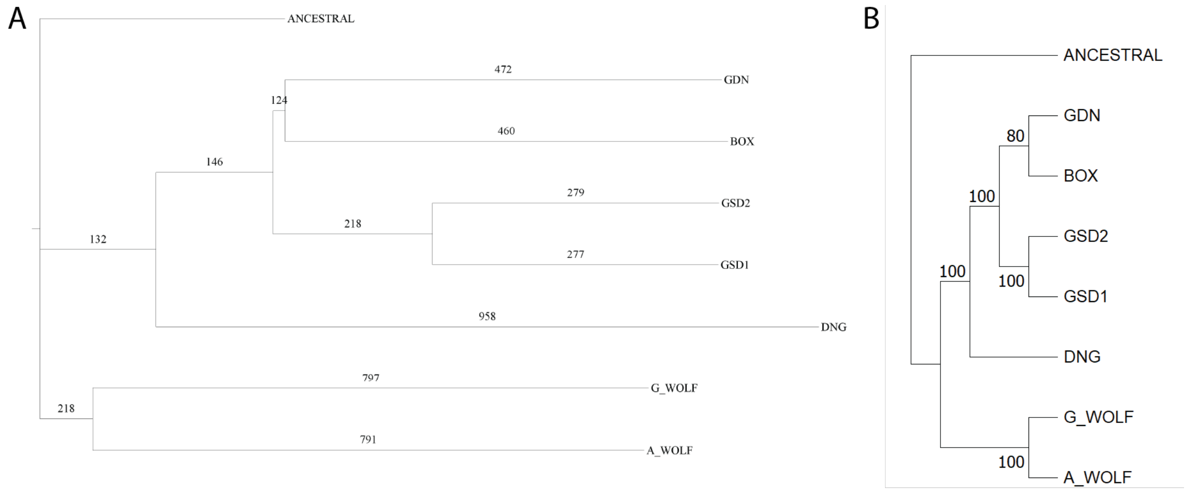


**Fig. S13: Phylogenetic Tree of samples utilizing dimorphic LINE-1 insertions.**

A phylogenetic tree was estimated using a distance matrix created from dimorphic LINE-1 loci. Trees were rooted on a theoretical ancestral genome for which all dimorphic LINE-1s are absent. The number of dimorphic LINE-1 variants is depicted on each branch (panel **A**). Bootstrap support from 1,000 runs is displayed on a phylogenetic tree with the same topology (panel **B**).

A

```
>G_WOLF chr21:11466940-11466940/rc
GCTGATAAGCTTTCTAATATAAAACAACAACAAGAAATACAAAT
AACAGGGCT

>GSD1 chr21:40000236-40006624
GCTGATAAGCTTTCTAATATAAAACAACAACAAGGAGGGGAGGA
GCAAGATGGCGGAAGAGTAGGGTCTCCAAATCACCTGTCTCCAC
CAAACTACCTAGAAAACCTTCAAATTATCCTGAAAATCTATGAA
TTCGGCCTGAGAATTAAAGAGAGACCAGCTGGAATGCA…TAGAA
GGGGAGGAGGGCGGGGGGTGGGAGTGAATGGGTGATGGGCACTG
GGTGTTATTCTGTATGTTAGTAAATTGAACACCAATTAAAAAAA
AAATAAAAAGAAAAAAAGAGAGAAAAATAAAAATAAAAAATAAA
ACAACAACAAGAAATACAAATAACAGGGCT
```

B

```
>G_WOLF chr26:26959991-26960043/rc
TGGAACTAAATATAAAACATACAATTAAAAAAAATCCTTGAAAA
AAATCCAAG

>GSD1 chr26:14961235-14967664
TGGAACTAAATATAAAACATACAATTAAAAAAAATCCTTGAGTC
CGGGGAGGAGCAAGATGGCGGAAGAGCAGGGTCTCCAAATCACC
TGTCTCCACCAAACTACCTAGAAAACCTTCAAATTATCCTGAAA
ATCTATGAATTCGGCCTGAGATTTAAAGAGAGACCAGCTGGAAT
GCTACAGTGAGAAGAGTTCGCGCTTCTATCAAGG…AGTGAATGG
GTGACGGGCACTGGGTGTTATTCTGTATGTTAGTAAATTGAACA
CCAATAAAAAATAAATAAATAAAAAAAAAAAAAAAAAATAAAATAA
AAAAATAAAAAAAATCCTTGAAAAAAATCCAAG
```

**Figure S14: Relaxed criteria identify interrupted poly(A) tails.**

Sequences corresponding to the filled and empty sites for two LINE-1 loci with interrupted poly(A) tails are shown. As in Figure 2, target site duplications are shown in orange in both the filled and empty site. LINE-1 sequence is shown in green; the three green dots indicate an artificial break in the sequence for display purposes. The inferred LINE-1 endonuclease cleavage site is underlined. The underlined sequence corresponds to the reverse-complement (top-strand) of the cleaved sequence. Interrupted candidate poly(A) tails are highlighted in yellow. Both examples are loci where GSD1 contains the filled site and G_WOLF contains the empty site. (**A**) A ~full-length dimorphic LINE-1 on chr21 is shown. No poly(A) tail was identified for this locus using the strict criteria because the sequence at the LINE-1/3′ TSD junction only ends with 5 adenosine nucleosides. (**B**) A ~full-length dimorphic LINE-1 on chr26 is shown. No poly(A) tail was identified for this locus using the strict criteria because the sequence at the LINE-1/3′ TSD junction only ends with 7 adenosine nucleosides.

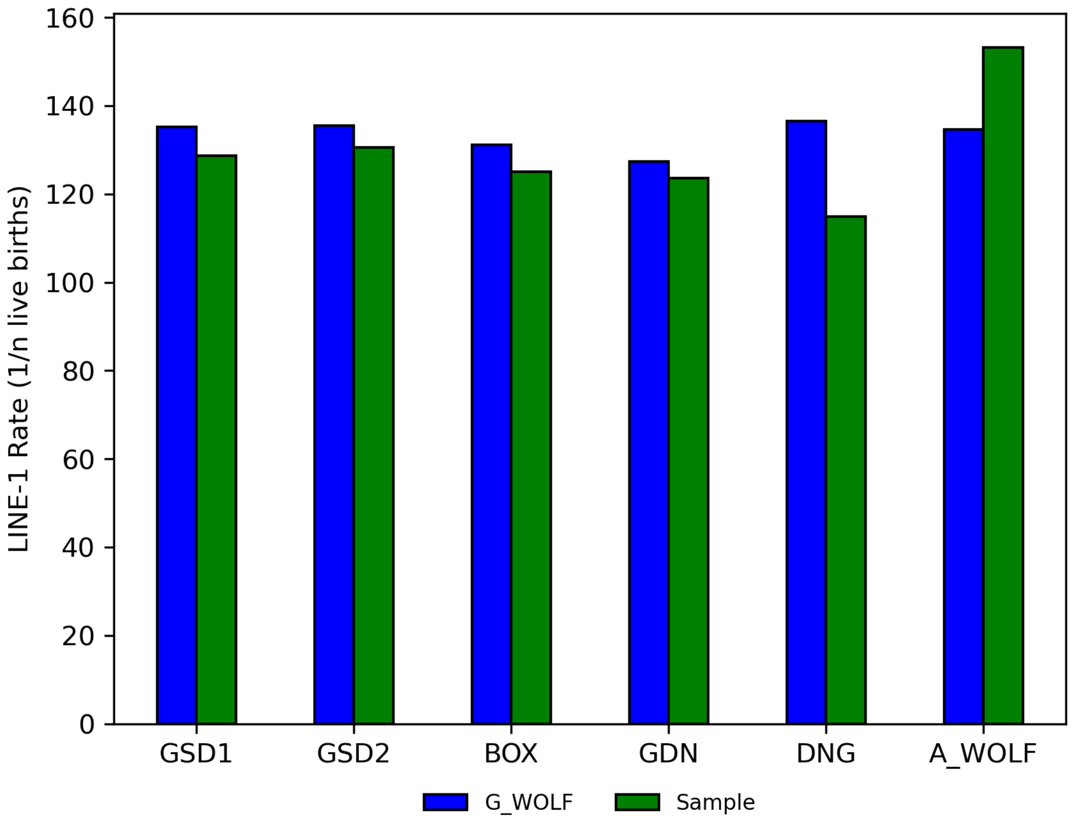


**Fig. S15: Side by side bar chart depicting estimated LINE-1 retrotransposition rates**

A bar plot depicts inferred SINEC insertion rates based on elements that are present in G_WOLF and not the queried sample (**Blue**) and in the sample but not G_WOLF (**Green**). Estimates are based only on autosomal data and assume a SNP mutation rate of $4.5 \times 10^{-9}$/bp/generation.

A

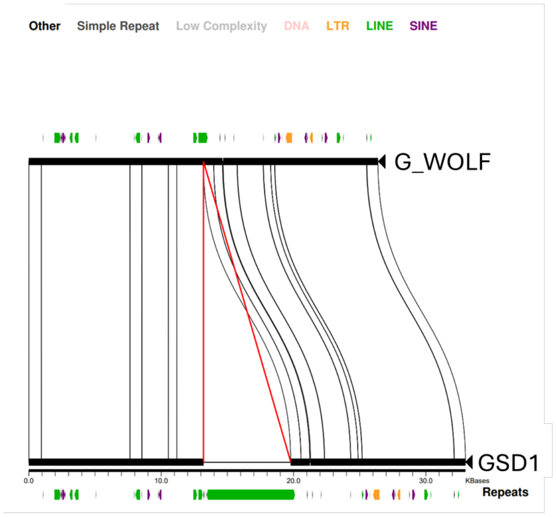


>G_WOLF chr1:92781149-92781199/rc
AAATCATGTTAGAGATTATAGGTAAATTTTAGACTCATCACAG
TTATAAGC

>GSD1, chr1:30077480-30084125
AAATCATGTTAGAGATTATAGGTAAGTTTTATTTTTTTTTTTT
TTTTTTTTTTTTTTTTTTTTTTTTTTTTTTTTTTTTTTTTTTA
ATGTTAAAAGGTCTCTTTATTTGAGATCAGACTTCCTAGGGC
AAGCTGTCTTTATAACTCTTCTTTATTTAATTCCCTTATTTGC
ACATTGAATAAATTAAAATGGCCATGGGAAAATTAAAATAATG
GTCCTTTTTTTTTTTTTTTTTTTTTTTTTGTTATCAACATTC
Gttttttgttttttttttttttattggtgttcaatttactaac
atacagaataatacccagtgcccgtcacccattcactcccacc
ccccgccctcctc......ccttgatagaagcgcgaactcttctcac
tgtagcattccagctggtctctctttaaatctcaggccgaatt
catagattttcaggataatttgaaggttttctaggtagtttgg
tggagacaggtgatttggagaccctgctcttccgccatcttgc
tcctcccccTTATAGGTAAGTTTTAGACTCATCACAGTTATAA
GC

B

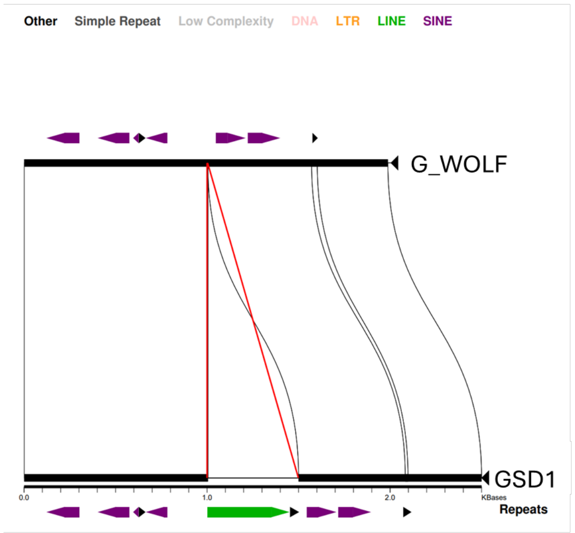


>chr1:90474713-90474764/rc
TCTTAGGGGAGGCAAAGTAGCTGGCTCAGAATGTGCTGGGAAG
GAGGCTATC

>chr1:32393866-32394413
TCTTAGGGGAGGCAAAGTAGCTGGCCcagcaatggccacgata
gccaaactgtggaaggagcctcggtgtccaacgaaagatgaat
ggataaagaagatgtggtttatgtatacaatggaatattactc
agctattagaaatgacaaatacccaccatttgcttcaacgtgg
atggaactggagggtattatgctgagtgaagtaagtcagtcgg
agaaggacaaacattatatgttctcattcatttggggaatata
aataatagtgaaagggaaaataagggaagggagaagaaatgtg
tgggaaatatcagaaaggagacagaacgtaaagactgctaac
tctgggaaacgaactaggggtggtagaaggggaggagggtggg
gggtgggagtgaatgggtgacgggcactgggtgttattctgta
tgatagtaaattgaacaccaataaaaaaataaattaaaaaaaa
aataaATAAAAAAAAAAAAAAAAAAAAAAAAAAAAAAAAAAAA
AAAAAAAAGAATGTGCTGGGAAGGAGGCTATC

**Fig. S16: LINE-1 insertions with target site deletions called by AGE**

Two LINE-1 insertions which AGE identified as possessing short target site deletions in the empty site were hand annotated. Both loci possess filled sites in GSD1 and empty sites in G_WOLF. LINE-1 sequence as called by RepeatMasker is depicted in green and lowercase, poly(A) tails are depicted in blue, TSDs are in gold, flanking sequence is in black, transductions are depicted in grey, and differences between GSD1 and G_WOLF are highlighted. Each comparison possesses a miropeats image of the locus. An insertion which AGE called with no TSD and a 1 bp deletion at the insertion site possesses a TSD which was not detected due to mutations between the left TSD, right TSD, and empty site (panel **A**). Another locus is truly missing a TSD in our dataset which may result from a target site deletion (panel **B**).
